# Lipid droplets in felid kidneys: prevalence and composition by lipidomics

**DOI:** 10.1101/2025.09.17.676785

**Authors:** R.A. Brociek, R. Alborough, M.A. Kotowska, A. Ferreira, S. Martinez-Jarquin, M. Walczak, F. Beaudoin, D.S. Gardner

## Abstract

An accepted and common phenotypic curiosity of Felidae is the presence of intracytoplasmic lipid droplets in renal proximal tubule epithelial cells (RPTEC), also frequently in urine (lipuria). Both outcomes are currently considered, and taught, as incidental – without obvious pathophysiological consequence. This contrasts markedly with clinical (human) medicine where lipid vacuoles in RPTEC are usually associated with metabolic or chronic disease, such as CKD. Despite domestic felids having a high incidence of CKD as they age, no study has fully characterised feline RPTEC lipid droplets in the context of CKD. Here, we first characterised the incidence of RPTEC lipid in domestic cat (with/without CKD or chronic interstitial nephritis) versus domestic dog and Scottish Wildcat, across a wide age range. Felids (domestic, wildcat) consistently had greater renal lipid content than dogs at all ages studied. Intracytoplasmic lipid extraction with chromatography, fatty acid characterisation and mass spectrometry-based lipidomics revealed unusual presence of a panoply of novel lipids found only in domestic cat: lipids were primarily modified (i.e. less polar) ether-soluble triacylglycerols including monoalkyldiacylglycerols (MADAGs) and other branched-chain fatty acids. We suggest common presence of such rare lipid species in tubular lipid droplets in domestic cat reflects an aspect of felid biology that parallels age-related disease prevalence, in particular, being associated with the aetiopathogenesis of chronic renal interstitial nephritis (CIN) – a hallmark of CKD in felids.

## Introduction

In humans and most other mammalian species, renal intracellular lipid deposition is an adverse finding associated with metabolic syndrome and/or obesity (Bobulescu 2010a). Ectopic lipid (i.e. lipid not esterified and stored in adipose tissue) can provoke and initiate renal cell damage through lipotoxic processes which may eventually result in CKD (Escasany, Izquierdo-Lahuerta, and Medina-Gómez 2018; Kang et al. 2014; Mitrofanova, Merscher, and Fornoni 2023). However, in the domestic cat, there is presence of a large quantity of intracytoplasmic lipid droplets within the renal tubular epithelial cells, and urine – lipuria, is accepted as an ‘incidental’ finding – that is, an observation with no known adverse outcome (Lucke 1968). Such early reports in the field of veterinary medicine are likely based on the fact that the droplets are so prevalent in cats, and do not appear associated with body weight or other co-morbidities (Modell 1933), and thus ‘incidental findings’ have become lingua franca in small animal veterinary medicine, in regard to the presence of lipid in felid kidneys.

In the domestic cat, renal lipid is mostly found in the proximal tubule epithelial cells (RPTEC) of the renal cortex (Lobban 1955; Foote and Grafflin 1938), although some deposits have been noted in the interstitium alongside macrophages (Martino-Costa et al. 2017). RPTEC lipid droplets or ‘lipid vacuoles’ have also been observed in wild or captive-wild cats such as lions and tigers, although the incidence is relatively low (Hewer, Matthews, and Malkin 1949; Newkirk et al. 2011). Nevertheless, lipuria remains a relatively common clinical pathological finding in wild cats (Hewer, Matthews, and Malkin 1949; D’Arcy 2018). Rarely are such phenotypes observed in domestic dogs or other mammals (Foote and Grafflin 1938). Advanced age is one factor known to increase the presence of renal lipids in both cats and in dogs (Macnider 1945); indeed, Quimby *et al* reported that 79% senior cats have renal lipid accumulation (Quimby et al. 2022). Remarkably, the same study reported 25% cats aged between 1 – 5 years also presented with renal lipid (Quimby et al. 2022), and at least some lipid droplets have been noted in kittens (Modell 1933).

Currently, no study, using advanced techniques to characterise lipids, has determined the composition of such intracytoplasmic RPTEC lipids in felids. An early study proposed that the lipid droplets were mainly triacylglycerides, phospholipids and cholesterol; (Bargmann et al. 1977). In plants (Chapman, Dyer, and Mullen 2012), animals such as the domestic dog (Maunsbach and Wirsén 1966) and humans (Olofsson et al. 2009), when such lipid droplets have been characterised then they are commonly described as dynamic organelles with neutral lipids in the central core, surrounded by a monolayer of amphipathic lipids (phospholipids and cholesterol), with a distinct proteomic profile (Bostrom et al. 2007; Ducharme and Bickel 2008). They are formed at microsomal (usually endoplasmic reticulum) membranes as primordial droplets with a diameter of 0.1–0.4 µm and increase in size by intracellular fusion. In skeletal muscle of athletes, intramuscular triacylglycerol-rich droplets are prevalent, despite being insulin-sensitive, as a cellular high-energy source (Stannard and Johnson 2004). The kidneys, particularly RPTEC, of all animals are highly metabolically active (equivalent to heart on a weight-specific basis) with high mitochondrial density (Else and Hulbert 1985). It is thus not beyond reason, that intracellular lipid droplets are related to cell metabolism and local microenvironment, although rarely have such phenomena been described. An additional contribution toward propensity for renal lipid accumulation is suggested by biological sex; for example, in male cats, renal lipid content increases after castration or when sexual activity is lost in older age (Lobban 1955). In contrast, Martino-Costa et al. (2017) concluded that interstitial lipid deposition affects male and female, neutered and non-neutered cats equally.

The precise composition of intracellular lipid droplets, when characterised in other species appears to be organ, tissue and cell-specific; for example, in liver, LD are described as comprising predominantly neutral, ether-soluble triacylglycerides, together with free or esterified sterols/cholesterol, surrounded by a phospholipid monolayer (Ducharme and Bickel 2008; Olofsson et al. 2009). Whereas, in one *in vitro* study using a kidney cell line (normal rat kidney, [NRK]) in which LD were induced by dosing with oleate, LDs also accumulated modified lipids such as monoalk(en)yl-diacylglycerols (MADAGS) (Wölk and Fedorova 2024; Bartz et al. 2007). In felids, little is known about the composition of the prevalent lipid droplets in kidney tissue. More extensive characterisation may help to understand whether such lipid droplets and the fatty acids that distinguish domestic cats from wild-cats and dogs, maybe causally linked to the mechanisms of CIN and CKD in cats.

Regardless, the presence of lipid droplets within cells of non-adipose tissue (i.e. ‘ectopic lipid’) is invariably associated with a chronic diseased/inflammatory state such as obesity and/or organ fibrosis, via mechanisms linked to cellular lipotoxicity (Thannickal et al. 2014; Wynn and Ramalingam 2012; Bobulescu 2010b; Sakuma et al. 2025). It is well known that both domestic (Lucke 1968; Lawson et al. 2015) and non-domestic felids (Newkirk et al. 2011) have a propensity towards, and indeed a high incidence, of chronic kidney disease. Yet, to date, in young, healthy cats, lipiduria and high concentrations of renal lipids remain an ‘incidental finding’. Any later causal association with feline CKD has not been proposed, despite the phenotype in cats being known for many years (Lobban 1955). Characterisation of lipids (‘lipidomics’) is a complex science and many lipids remain unclassified. Esterified triacylglyceride is accepted as the common storage form of lipid (Bobulescu 2010a); phospholipids (e.g. PC, PE) are structural components of biological membranes (Pan, Hulbert, and Storlien 1994); ‘00s of individual fatty acids exist in water-soluble, non-esterified form and participate in intracellular energy cycles’ (Stich and Berlan 2004). Some lipid classes are considered toxic when accumulated in various organs – diacylglycerides, ceramides and cholesterol (Weinberg 2006; Sakuma et al. 2025). Inter-organ exchange of lipid is a normal metabolic process and without consequence, unless lipids are deposited in the interstitium (i.e. tubulointerstitium in kidneys) where lipotoxicity elicits inflammatory reactions that underpin chronic interstitial nephritis/inflammation (CIN), a hallmark of CKD in felids (Martino-Costa et al. 2017; Mitrofanova, Merscher, and Fornoni 2023). And yet, no studies in companion animals have linked the common findings of renal intracellular lipid with CIN and CKD. Hence, further investigation of this phenomenon in cats is warranted.

In this study, the renal lipid content of domestic cats (DC), domestic dogs (DD; as comparator domestic species not prone to CKD) and Scottish Wildcats (SW; as a felid, but eating unprocessed food) has been characterised and described in pathological tissue from otherwise healthy (i.e. did not die or was euthanised for renal causes) or from cats with histopathologically diagnosed renal disease (e.g. diagnosed CKD). We hypothesise that 1) domestic cat kidneys contain more lipid than domestic dogs or Scottish Wildcats, 2) the lipid content of the domestic cat’s kidney is increased during renal disease and 3) the composition of RPTEC lipids in domestic cat kidney is unique and underpins propensity to CKD in domestic cats.

## Materials and Methods

### Sample collection and preparation

Initial studies (*ca*.2018-2020) characterising lipid droplets (e.g. Oil-Red O) in domestic animals without renal disease used kidney tissue from n=21 domestic cats, n=22 dogs and n=8 Scottish Wildcats. A further n=14 domestic cats with histopathologically defined chronic interstitial nephritis (CIN) were also obtained. All samples were from Veterinary Pathology, School of Veterinary Medicine and Science. Ethical approval for use of pathological tissue was given by the Committee for Animal Research and Ethics (CARE), University of Nottingham on a number of occasions (REC: 3256 201020; 3900 230809; 4055 240124). Further samples for chromatographic and mass-spectrometry studies were obtained from similar sources (e.g. Veterinary Pathology; domestic cats (n=21), dogs (n=8)). Samples were representative of male and female cats/dogs, drawn from a wide age range and body condition and were predominantly neutered. In addition, opportunistic samples from non-domestic feral wildcats were obtained from National Museums Scotland (Scottish Wildcats [Felis silvestris] (n=21) and captive zoo wildcats (n=3)). A variety of measurements determined age and species (full wildcat vs domestic cat hybrid). All kidneys were stored between -20°C and -80°C upon collection, and studies described herein were conducted between 1 month and 2 years between collection and processing.

### Histological evaluation and quantification of renal lipids

haematoxylin and eosin stained FFPE kidney tissue sections (5 µm) were used to visualise gross pathology by light-microscopy (Nikon *i*50/80; Nikon Digital Sight DS-U1). Frozen kidney tissue sections (10µm) from n = 21 domestic cats, n = 22 dogs and n = 8 Scottish wildcats were prepared onto polylysine slides at - 20°C for staining of tissue lipid using Oil Red O (ORO). In brief, using n=3 different samples of adipose tissue as a positive control, sections were immersed in 1% ORO in 70% industrial methylated spirit (IMS) for 30 mins, further rinsed in 30% IMS, washed in running tap water, rinsed in distilled water, then counterstained with haematoxylin (30 secs). Finally, after washing in tap water, blued sections were aqueous mounted and a coverslip applied. All sections were pseudoanonymised (e.g. N85-2345) before ORO quantification: n=10 regions of interest in the cortex were assessed per section (random selection using a zig-zag construct) and images saved. Image J was used to quantify greyscale images of percentage ORO positive areas (ORO^+ve^ – as proportion of total area with tissue). Imaging parameters (hue, saturation and brightness) were initially optimised then fixed and applied to all sections. An average ORO^+ve^ area per slide was calculated.

### Extraction of renal intracytoplasmic droplets

First a lipid extract was isolated from kidney tissue using the sucrose cushion method of lipid extraction by ultracentrifugation (Figure S1a-m). In brief, a 1 litre stock solution of isolation buffer (50 mM HEPES (4-(2-hydroxyethyl)-1-piperazineethanesulfonic acid), 10 mM potassium chloride, 62.5 mM potassium acetate, 5 mM EGTA (ethylene-bis(oxyethylenenitrilo)tetracetic acid), 5 mM DTT (1,4-dithiothreitol), 1 mM magnesium chloride and 5 mM EDTA; pH = 7.5) was prepared. Approximately 1g of frozen renocortical tissue was homogenised on ice in 2 ml of 0.6M protease-free sucrose (with 1% BSA) extraction buffer, using a gentleMACS™ tissue dissociator. The homogenate was then centrifuged at 3222 rpm (2000*g*) at 4°C for 5 minutes and 2ml of the supernatant overlayed on 1ml 0.6M protease-free sucrose extraction buffer. Finally, 2mls 0.25M protease-free sucrose (with 1% BSA) extraction buffer was overlayed on top to create a discontinuous sucrose cushion. The layered sample (in a 5ml ultracentrifuge tube) was then ultracentrifuged at 100,000*g*, 4°C for 1 hour (Hitachi CP80NX, P55ST2 rotor). After, the tube with a turbid layer of cytoplasmic lipid at the top was frozen overnight at -20°C. The top 1cm of a 5ml ultracentrifugation tube was removed, being representative of total lipid extract (TLE; Figure S1i). To 1 ml of the TLE, 5 ml of 2:1 (v/v) chloroform:methanol was added in a 15ml glass centrifuge tube, vortexed twice (15 secs) before adding 1ml of 1% NaCl. Further vortexing ensured a homogenous lipid solution (soln. A) which was centrifuged (1000*g* for 2 mins). The lower organic phase, comprising the lipids of interest, was transferred into a new glass vial (soln. B). 2 ml of 100% chloroform was added to soln. A, vortexed twice for 15 seconds and centrifuged again at 1000*g* for 2 mins. The lower organic layer was again transferred and pooled with soln B, before drying to completeness under nitrogen gas, re-suspended in 200µl 100% chloroform and stored in vials at -80°C.

### Characterising and describing the lipid extract

#### High Performance Thin layer chromatography (HPTLC)

The separation and identification of TLE lipids was performed using HPTLC. 5-10μL of the lipid standard mix containing 1,3-Diolein, 1,2-Dioleoyl-rac-glycerol, Glyceryl trioleate, Monoolein (1787-1AMP, Merck, UK) and 10μL of TLE were applied on a HPTLC silica gel 60 F254 (20x10cm) glass plate (Merck, USA) using a Linomat V (Camag, Switzerland) sample applicator. To maximize chromatographic resolution and facilitate superior analyte separation, the plate was developed twice in a chromatographic chamber (Camag, Switzerland) saturated with a mobile phase of hexane:diethyl ether:acetic acid (68:12:0.4 (v/v/v)). Once dry, the plate was in a solution containing 10mg Primuline dissolved in 200ml acetone:water (80:20 (v/v)) using The CAMAG® Chromatogram Immersion Device 3 (Camag, Switzerland) set at immersion speed ’1’ and duration setting ’6’. The plate was scanned RT White, at 254nm and 366nm using TLC Visualiser 2 (CAMAG, Switzerland). The bands of interest were marked in pencil (e.g. unidentified band or ‘UB’; see Figure 2) and scraped for further analysis. Image profiles were generated, and data was processed using VisionCATS version 3.2 software (Camag, Switzerland).

#### Isolation of specific lipids from HPTLC bands of interest

for analysis of known or unknown lipids in specific bands on the TLC, then bands were scraped using a flat-end spatula and transferred to a 5ml glass vial. In negative samples (i.e. no visible ‘UB’), a similar area was also scraped, for comparison. 1ml of 100% chloroform was added and gently agitated before heating at 35°C in a waterbath for 20 minutes. Any remaining chloroform was transferred to a new vial (‘Vial A’), taking care to not include silica particles. Silica particles had a further 1ml 100% chloroform added, re-agitated for 15 secs, and remaining chloroform, but not silica, pooled together with ‘Vial A’ and frozen at -20°C (see Figure S1).

### FAME lipid extraction and separation

To the extracted lipid suspension, 0.7mL 10M KOH and 5.3mL methanol were added. The sample was heated at 55°C for 90min. After cooling, 0.58mL 12M H_2_S0_4_ was added and further incubated at 55°C for 90min. After cooling, 3mL hexane was added, mixed and centrifuged. The upper hexane layer was removed, concentrated by drying under nitrogen and reconstituted in 400µL hexane before storing at -30°C until analysis by GC-MS analysis. ***Gas chromatography-flame ionisation detection (GC-FID) of FAMEs*:** The fatty acid methyl esters (1µl) were injected (split ratio 50:1) into a gas chromatograph (GC) (Trace 1300, Thermo Fisher Scientific™) coupled with mass spectrometer (MS) (ISQ 7000, Thermo Fisher Scientific™). Separation of fatty acid methyl esters was performed with a Varian CP-Sil 88 (100m length, 0.25mm diameter, 0.20um film thickness, Agilent) capillary column with helium as carrier gas. Oven temperature (ramp up at 4°C/minute, from 140°C (hold for 5 minutes) to 240°C (hold for 10 minutes) and MS injector and transfer line temperature (260°C and 250°C, respectively) were preprogrammed. The ion source temperature was set to 200°C. Characterization and identification of FAMEs was performed in scan mode. Quantification was completed by selective ion monitoring (SIM) mode of the most intense fragments. Data acquisition and processing were performed with Chromeleon (version 7.0, Thermo Fisher Scientific™). Quantification was based on external calibration with C19:0 as an internal standard.

### Liquid Chromatography-Mass Spectrometry (LC-MS)

The lipidomics liquid chromatography (LC) method was based on that previously described (Whitby et al. 2023). For the LC an Ultimate 3000 HPLC (Thermo Scientific) was used in reverse-mode with an ACE Excel 2 SuperC18 column (50 2.1 mm; 2 m particle size; Advanced Chromatography Technologies, Aberdeen, UK), held at 50°C. The starting mobile phase was 30% B (10% water, 0.1% ammonium acetate, 10% acetonitrile, 80% isopropanol) and 70% A (60% water, 0.1% ammonium acetate, 40% acetonitrile) at a flow rate of 400 µL/min, increased to 35% B by 1 min, then to 100% B by 7 min. The flow rate was then increased to 500µL/min by 11 min. The proportion of mobile phase B decreased to 20% by 12 min, equilibrating for 3 min. For MS, the QExactive Plus Orbitrap (Thermo Fisher Scientific, Hemel Hempstead, UK) was used for LC-MS simultaneous ESI+ and ESI-modes. The probe and capillary temperatures were maintained at 412.5 and 256.25 ºC, respectively. The following settings were used: sheath gas 47.5, auxiliary gas 11.25, and sweep gas 2.25, AGC target 3. The spray voltage was set to +4.0 kV or –4.0 kV. All files were acquired to get a full scan (70,000 resolution). Then data-dependent tandem MS/MS (ddMS^2^) spectra were produced on the five most intense ions at any one time at a resolution of 17,500 20, 30 and 40, isolation window 1.0 m/z, intensity threshold 1.6 10x^5^, dynamic exclusion 8s.

For later experiments, the LC-MS conditions were modified to achieve greater resolution and specificity: mobile phase A and B remained the same but after the starting ratio of 70% A and 30% B at a flow rate of 400 µL/min was increased to 35% B by 1 min, to 100% B by 7 min with the flow rate increased to 500 µL/min by 11 min (600 µL/min by 12 min). Finally, the flow and gradient became 400 µL/min and 30% of B by 13 min, equilibrating at this level for 6 min for a total run time of 19 min. For the Mass Spectrometry analysis, a Lumos Fusion Trihybrid instrument (Thermo Fisher Scientific, Hemel Hempstead, UK) was used. To enhance MS^2^ quality, data were acquired separately in positive and negative modes. The spray voltage was 3.5 kV for the positive mode and 2.5 kV for the negative mode. The sheath gas was 50, aux gas was 10, and sweep gas was 1, all in arbitrary units. Ion transfer and vaporiser temperatures were 325 and 350, respectively. A predesigned workflow of General lipid profiling MS^2^ was adapted. The MS1 resolution was 120,000 with a scan range of 250-1500. Data-dependent MS^2^ was performed with an isolation window of 1.5 m/z. Assisted HCD Collision Energy was used with values of 15, 30, and 45 eV. Data acquisition was carried out at a resolution of 15,000.

### In situ lipidomics of frozen kidney sections using OrbiSIMS

Orbitrap secondary ion mass spectrometry (OrbiSIMS) enables label-free imaging of tissues (Starr et al. 2022) and cataloguing of complex biological sample chemistry (Kotowska et al. 2023). Here, 10 µM sections of frozen kidney cortex from domestic cat (n=2), Scottish wildcat (n=2) and captive wildcat (n=2) were obtained using a cryotome (Leica Biosystems) at -20°C, adhered to glass polylysine slides and stored at -20°C until required. Cryotome samples were transported on ice to the school of Pharmacy (University of Nottingham) for OrbiSIMS analysis. Slides were transferred into a Leica VCM bath under liquid nitrogen, attached to a Leica cryogenic block. Blocks were introduced into an airlock on the cryo-sample holder via a Leica vacuum transfer system. Analysis was carried out at −170°C using a closed-loop liquid nitrogen pumping system (IONTOF GmbH). For acquisition of OrbiSIMS images, a 20 keV Ar_3064+_ analysis beam of 2 µm diameter with duty cycle set to 27% was used as primary beam, current at 24 pA. Obtained images represented an area of 400 µm × 400 µm using random rastor mode. Pixel size was 4 µm, thus total number of pixels was 100 × 100 over a cycle time of 200 μs. Argon gas flooding was in operation in order to aid charge compensation, with pressure in the main chamber maintained at 9.0 × 10^−7^ bar. Spectra were collected in negative polarity, over a mass:charge (m/z) range of 75 – 1125. Injection time was 500 ms. Mass-resolving power was 240,000 at m/z 200.

### Data analysis and presentation

Data was collected using a multitude of analytical equipment and analysis software, as indicated in the main text. Data is primarily presented as raw data to show distinct differences between individuals. VisionCats (CAMAG) and Freestyle (Thermo Fisher Scientific) were used to visualise data. Lipid identification was completed using a combination of Lipidsearch 5.1(Thermo Fisher Scientific) and LIPID MAPS®. Quantitative data was processed and presented using GraphPad Prism v10.3.0 (GraphPad Software Inc., California, USA). Unpaired t-tests were used to compare OrbiSIMS groups.

### Data availability

Any data relevant to this manuscript are available from the authors on reasonable request.

## Results

### Oil-Red-O staining of kidney tissue

The proportion of kidney tissue positively stained with Oil Red O (ORO^+ve^) was similar between domestic cats and Scottish wildcats, but each had significantly greater lipid content than domestic dog (domestic cat, 18.48% [9.069%, 25.66%]; Scottish wildcat, 14.09% [6.756%, 22.90%]; dog, 1.002% [0.0917%, 13.63%], median [IQR]; *P* = 0.018 (Figure 1A,C). As a positive control and for reference, sections (n=3) of adipose tissue were 54.76% [53.18%, 63.92%] ORO^+ve^, as expected (Figure 1A). At all ages available, domestic cat had greater lipid content than dog (Figure 1B), even in the few samples available ≤2 years of age. The intercept of the lines of best fit for cats and dogs was significantly different by linear regression (P = <0.001). Over time, in both species, it was apparent that the lipid content of kidneys gradually declined rather than increased (Figure 1B), likely reflecting the age-related loss of renal proximal tubule epithelial cells (RPTECs) which primarily harbour lipid droplets in feline kidneys. Next, it was important to characterise and describe the identity of the prevalent cytoplasmic lipids in cat (*cf*. dog), and a number of lipidomic approaches were used to do this.

**Figure 1.**
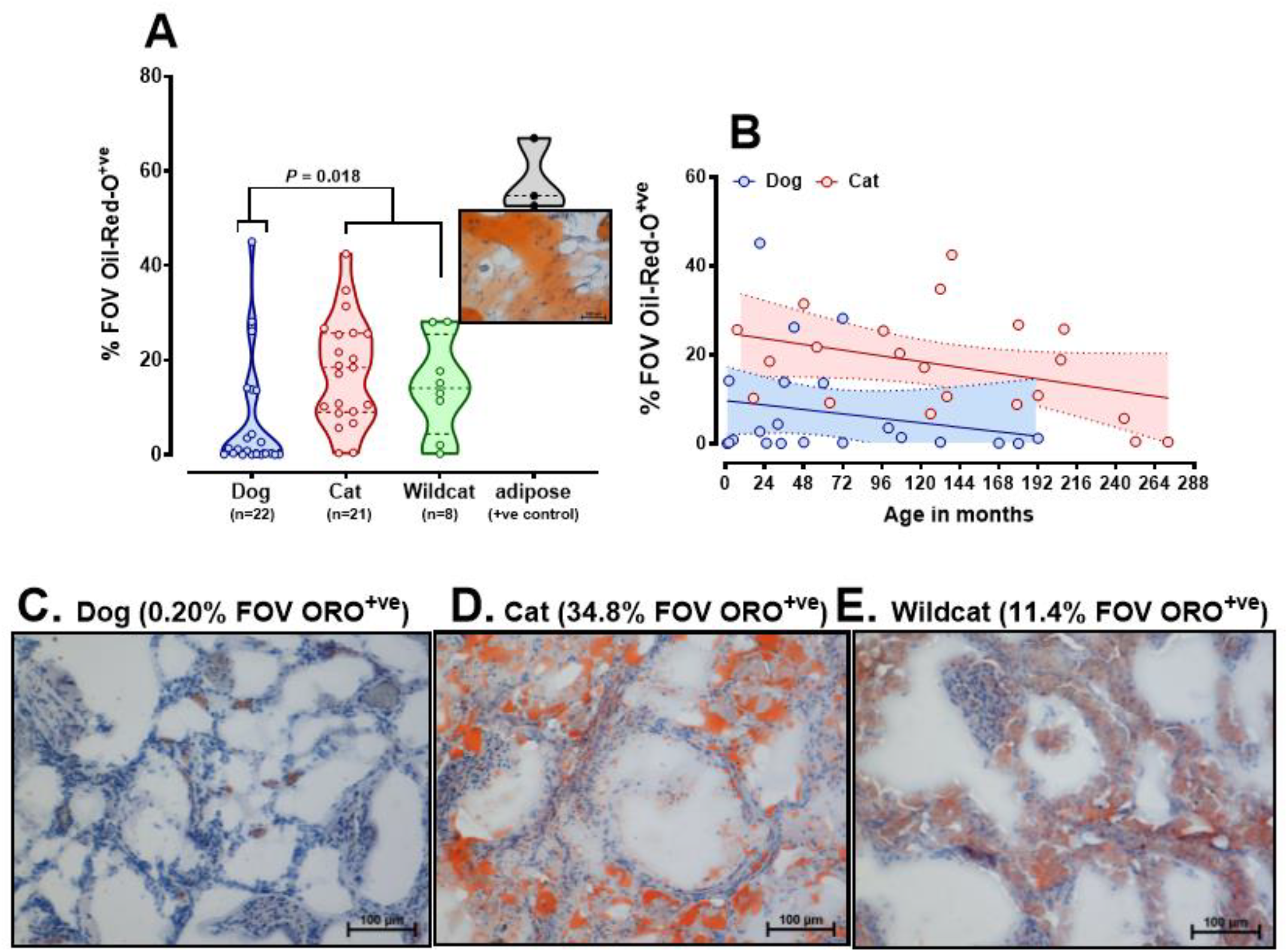
Oil-Red-O (ORO) staining of lipid in domestic cat, Scottish wildcat and dog kidney sections. **(A)** Violin plot of individual samples (represented as circles) of domestic cat, Scottish wildcat and dog kidney sections, stained with Oil Red O, using adipose tissue as a positive control. Data are average of ten random fields of view stained positively with Oil Red O per slide. The full range of data is depicted within the violin plot, dashed lines representing the lower and upper quartiles, with median. Lipid content of cat and wildcat kidneys were similar, each being significantly greater than dogs (*P* = 0.018, NP-ANOVA). **(B)** Scatter graph of percent ORO^+ve^ versus age of individual. Solid lines are lines of best fit, dotted lines indicate 95% confidence interval around the mean for age. **(C-E)** Representative images of ORO stained kidney sections from **(C)** dog, **(D)** domestic cat and **(E)** Scottish wildcat. Images were taken with a Nikon Digital Sight DS-U1 camera, Nikon i80 light microscope, 20x objective.

### High Performance Thin Layer Chromatography (HPTLC)

Total lipid extract (TLE) separation of major lipid classes by HPTLC of n=24 domestic cats (DC), n=12 domestic dogs, n=6 Scottish wildcats (SW) and n=3 captive wildcats (CW) indicated an unusual, unidentified lipid class (hereafter referred to as ‘unidentified band’ or UB) in the majority of DC (i.e. 23 of 24 domestic cats). The lipid was less polar than triacylglycerols (TAG), the predominant lipid present in most samples, and was more polar than cholesterol esters (CE; see Figure 2A-D). Further analyses confirmed the UB to be present in all adult DC kidney samples, whether healthy (n=12) or DC with a previous diagnosis of chronic interstitial nephritis (CIN; n=12), the defining histopathophysiological feature of CKD in DC. No canine or captive wildcat (CW; n = 3 [n = 2 snow leopards and n = 1 tiger]; Figure 2B) exhibited a UB. It was occasionally observed, however, in adult Scottish wildcats (see Figure 2C). Indeed, further analyses of TLE by HPTLC from a further n=18 domestic cats, n=8 domestic dogs, n=21 Scottish wildcats and n=3 captive wildcats confirmed previous observations. Most TLEs contained common lipids present in the lipid reference, e.g. monoalkylglycerol (MAG), 1,3-diacylglycerol and 2,3-diacylglycerol (DAG), free fatty acids (FA) and triacyclglycerol (TAG). The unidentified band present with an Rf value

**Figure 2.**
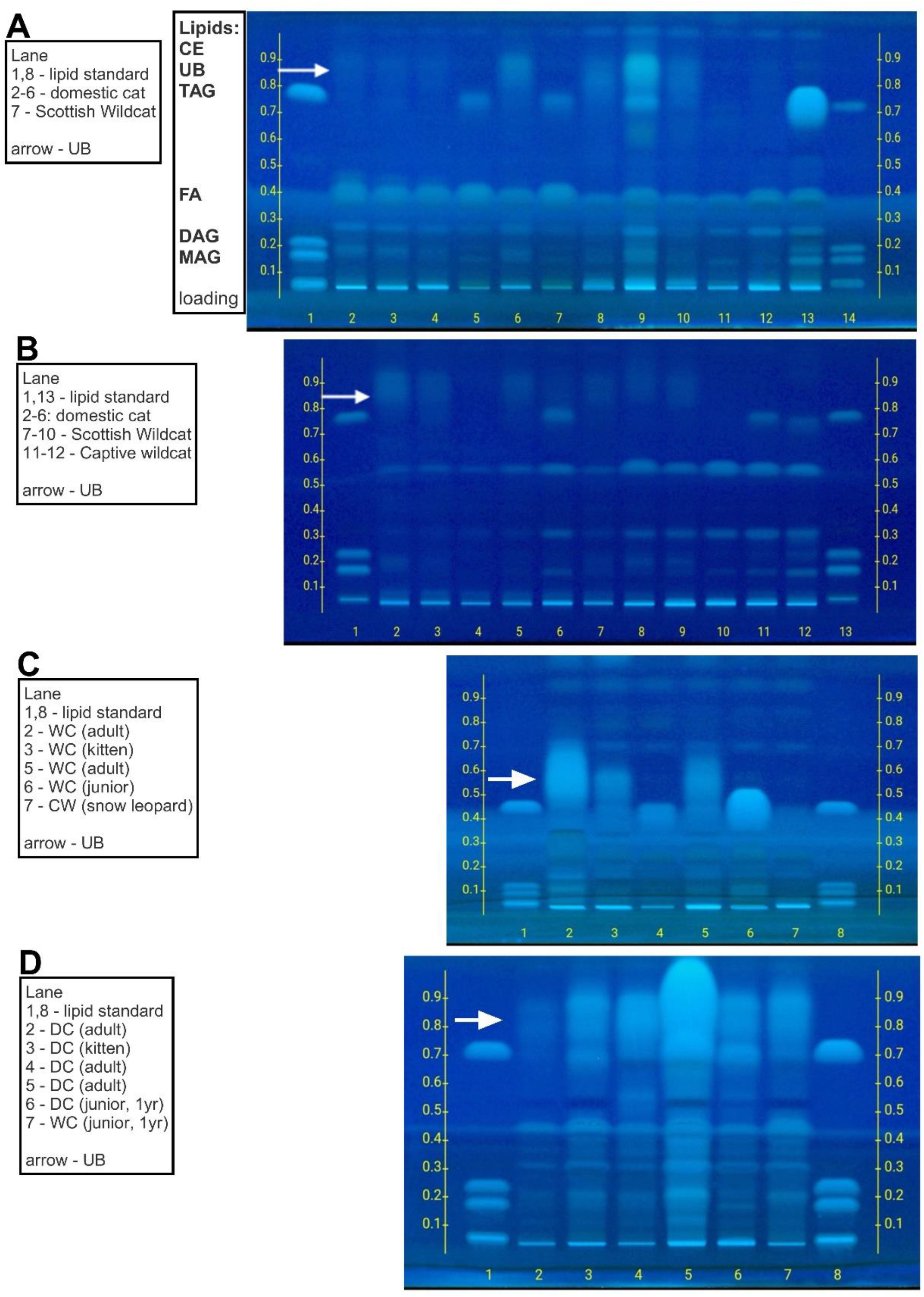
High Performance Thin Layer Chromatography (HPTLC) separation of major lipid classes in domestic cat (DC), Scottish wildcat (SW), Captive Wildcat (CW) and dog kidney after extraction and isolation of intracytoplasmic lipid droplets. **A-D:** High Performance Thin Layer Chromatography (HPTLC) separation of major lipid classes in domestic cat (DC), Scottish wildcat (SW), Captive Wildcat (CW) and dog kidney. Each lane represents a separate sample with lipid standard (containing mono, di and triglyceride; [MAG, DAG and TAG]) run first and last on each plate. Further detail of each individual is given in Supplementary Information, item 2. Arrow indicates relative position of unidentified band (UB) on each plate. All unknown samples were loaded at 10µL, all lipid standards at 5µL. TLC carrier phase was hexane:diethyl ether:acetic acid (68:12:0.4 (v/v/v)).

0.9 – 1.0, between TAG and CE often substituted to some extent for TAG, but this was dependant on age. For example, in kittens often only the TAG band was present, whereas in elderly individuals then the TAG band was diminished but the UB prominent. In captive wildcats, only the TAG band was prominent (Figure 2C-D). Evaluating this phenomena semi-quantitively using HPTLC chromatograms, then the ratio of TAG:UB varied consistently with age: in kittens, TAG:UB was 22.6:20.0% UB versus adult domestic cats where the ratio was 6.4:25.3%. Scottish wildcats were similar; kittens, 24.1:14.0% versus 7.8:21.2% in adults. Dogs and captive zoo wildcats were consistently ∼ 15.1 - 16.1% TAG, but negative for the UB. Thus, extraction of intracytoplasmic lipid from various felids with dogs as a domestic comparator/reference species reveals the virtually exclusive presence of an unidentified lipid band in domestic cats. The band is likely similar to TAG but less polar, suggesting modification to the fatty acyl chains of TAG. The presence of such ‘modified TAGs’ appears less often in kittens or other young felids, despite lipid droplets being evident at a histological level at this age (Figure 1B), but becomes more prevalent with age, often substituting/replacing TAG at this time, consistent with the UB being ‘modified TAG’. We therefore further characterised the UB in terms of fatty acid composition and by using mass-spec based lipidomics.

### Liquid Chromatography Mass Spectrometry of total lipid extract (TLE) from kidneys

Total lipid extracts from n=19 individuals (n=9 domestic cats, n=4 domestic dogs and n=6 Scottish wildcats) were assessed by LC-MS, with annotation of lipid classes using Lipidsearch™. A total of n = 2353 different lipids were identified. As expected, Scottish wildcat and domestic cat had greater (as proportion of total) TAG relative to dog (DC, 71.6; SW, 71.8 vs. Dog, 39.4%; Figure 3A). Phospholipids were the other major contributors to the lipids in TLE between species; Figure 3B). For example, the TLE from dog comprised 31.5% phospholipid (versus DC, 16.7% and SW, 12.2%), primarily comprised of phosphotidylcholine (59.3%, coloured purple in Figure 3B) and lysophosphatidylcholine (14.3%, blue). DC and SW were broadly similar in overall profile, although DC had greater phosphatidylcholine (66.8%), whilst SW had greater phosphoglycerol (9.8%) relative to DC (coloured red in Figure 3B). Thus, the type of lipids in companion animal kidneys are highly neutral; after TAG and phospholipids, no other lipid contributed >3%. Characterising the types of TGs (which were numerous, Figure 3C) that make up the TAG in the lipid droplets extracted from kidneys, revealed some interesting differences between dog and felids: in dogs, far fewer species were present, dominated by three individual TGs (50:2, 50:3, 50:1), whereas felids (domestic cat and SW) had far greater number of detectable species in total cf. dogs, but were broadly similar between DC and SW (Figure 3C).

**Figure 3.**
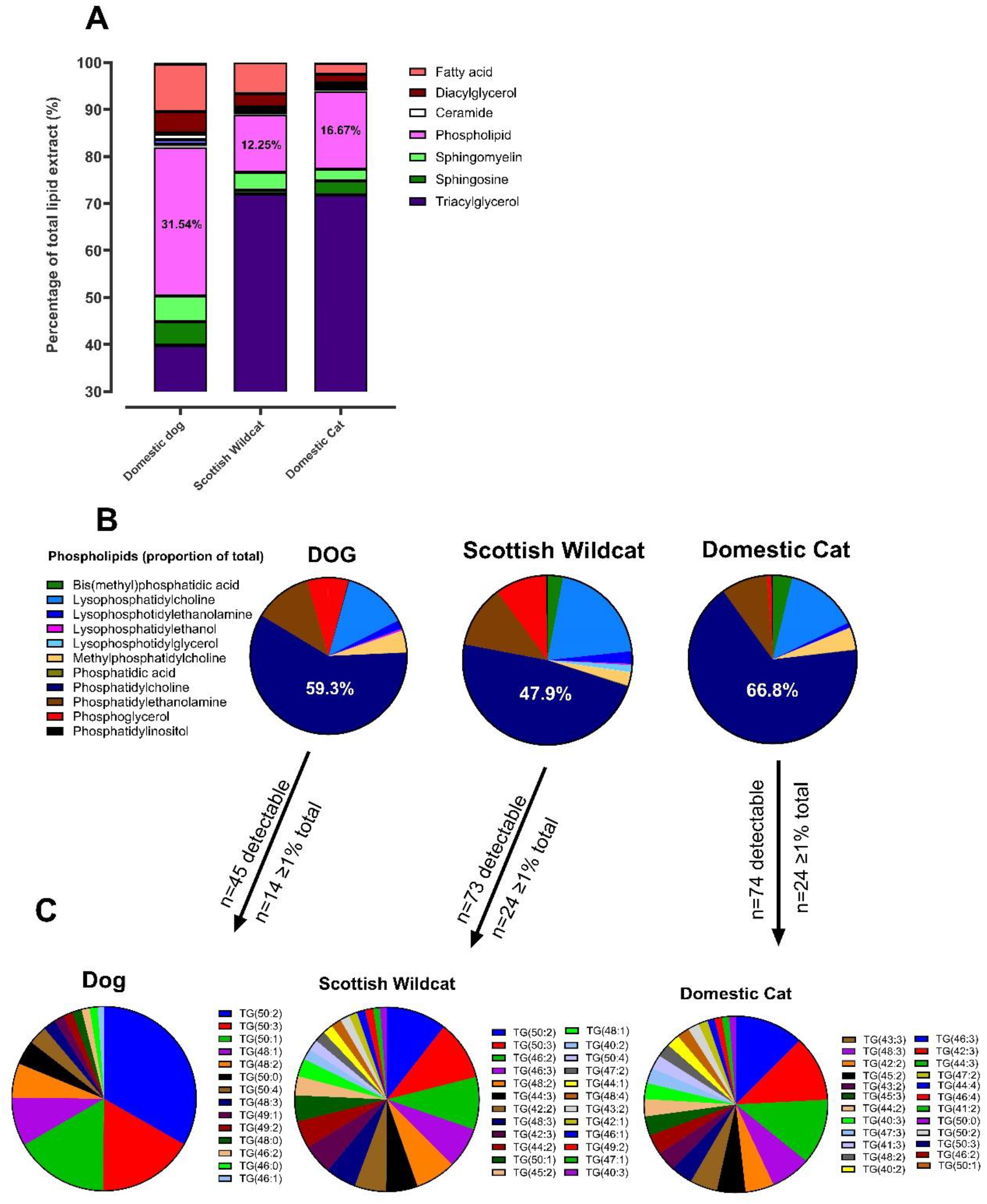
LC-MS analysis of lipids in dog, Scottish wildcat and domestic cat kidneys. Percentage composition of lipids in of dog (n=4), Scottish wildcat (n=6) and domestic cat (n=9) total lipid extract by percentage of total, as determined by LC-MS. Average group (as determined by Lipidsearch) composition of dog, SW and DC **(B)** phospholipids and **(C)** triacylglycerols >1% of total.

Comparing the 15 most abundant lipids extracted from each species kidney, then DC was almost exclusively comprised of shorter, mono-unsaturated fatty acids such as (14:1_14:1_16:1) and (14:1_16:1_16:1) which were also present in Scottish wildcat but undetectable in domestic dog (Figure 4A,B). In the latter, combinations of common saturated fatty acids were dominant such as palmitic (16:0), oleic (18:1) and linoleic (18:2) acid (e.g. TG(16:0_16:1_18:1) and TG(16:0_16:0_18:2), Figure 4A). In Scottish wildcats, the top 15 lipids were also present in DC, but at much lower levels (e.g. only one of the top 15 also appeared in DC; (TG(14:0_14:1_16:1)) and were virtually absent from domestic dog (Figure 4C). Thus, felids have more TAG in their kidney cortex than dogs, with a much greater phosphatidylcholine to total TAG ratio, and far greater TG species – for felids, these TGs appear to have an unusually high proportion of short-medium chain, mono-unsaturated FA. Such differences could, partially, affect polarity and may explain the polarity shift from TAG to UB, by TLC. We therefore chose to specifically isolate (i.e. by scraping and extracting the UB from the TLC plate) and characterise the individual lipids in the UB (cf. total lipid extract) by mass-spec lipidomics and derivation of fatty acid methyl esters.

**Figure 4.**
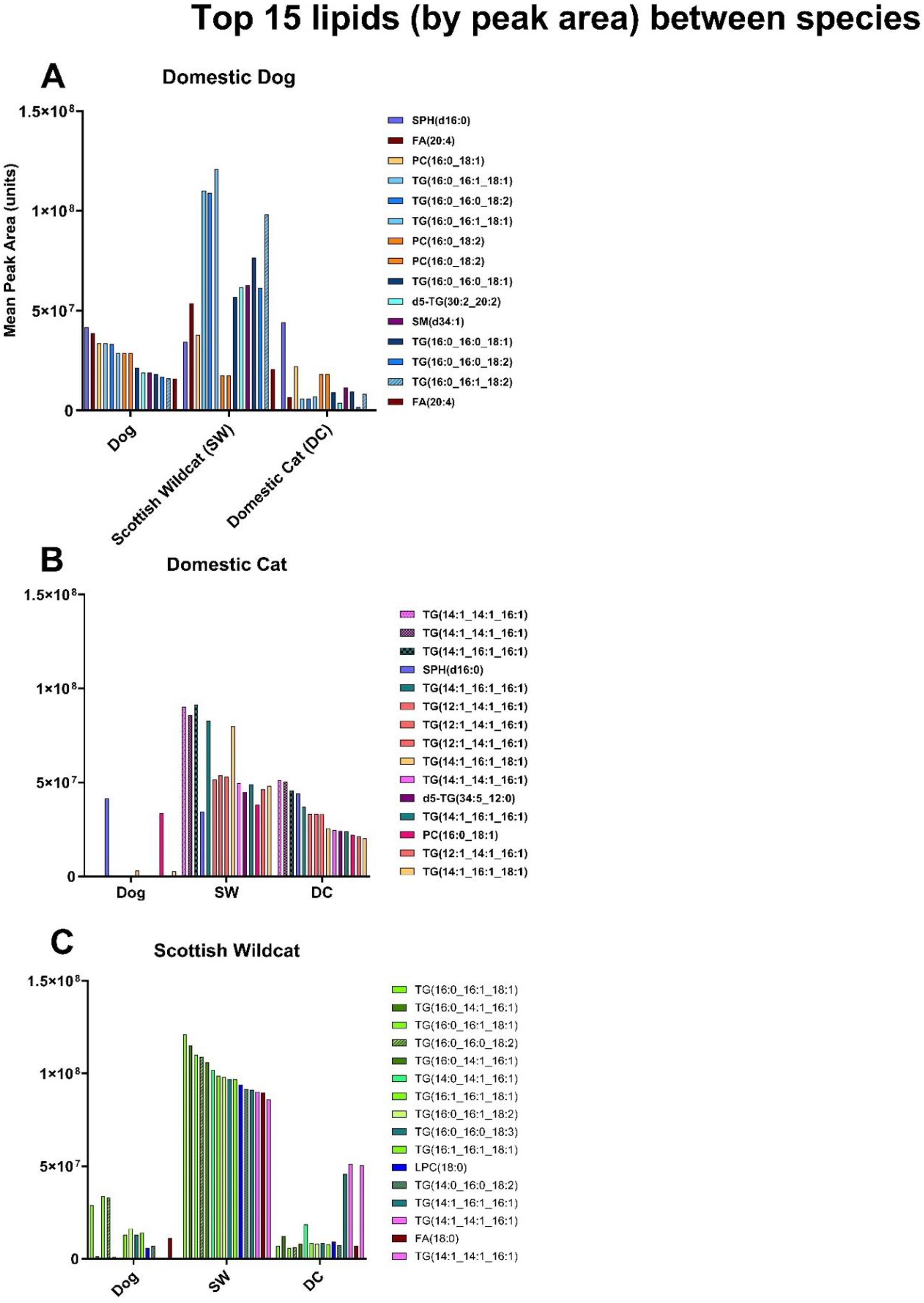
Top 15 lipids extracted from kidneys (Dog, Domestic cat, Scottish Wildcat) Top 15 lipids detected in renal cortical tissue of **(A)** domestic dog, **(B)** domestic cat and **(C)** Scottish wildcat. Bars represent mean peak area for each lipid across all individuals of that species. Each colour corresponds to a specific lipid identity, and is consistent across A-C. While some lipids are shared between species, the top 15 lipids from each species are largely dominated by different lipids.

### Fatty acid methyl esters (FAMEs) in the UB

Individual lanes on TLCs from felids were categorised as either positive (UB^+ve^) or negative (UB^-ve^) for visual presence of the unidentified band (see example, Figure 5a). The area was visualised, scraped and extracted appropriately for analysis of FAME content (Figure 5b-d). A similar process was adopted for various TAG standards, to confirm recovery of expected fatty acids (e.g. stearic, C16:0 and palmitic, C18:0). In most samples from total lipid extracts and the TAG band, the majority fatty acids were, as expected, C16:0 and C18:0 series (Figure 5b,d). Notably, in the UB^+ve^ band – largely exclusive to DC – C12:0 (lauric acid) and C14:0 (myristic acid) were common, whereas in dogs it was absent (Figure 5c). Nevertheless, in the protocol for methylating FAMES, an ester bond is cleaved in exchange for an alcohol functional group. Thus, it is perhaps unsurprising that further FAME analysis did not reveal any marked/notable differences between the species, beyond a preponderance of short-to-medium chain FA in DC (such as caproic, [C6:0]; lauric [C12:0] and myristic [C14:0], Figure 5e,f). FAMES, which derivatises lipids with ester bonds, would not report difference in lipids with potential ether bonds, which would reduce their polarity due to addition of a hydroxyl group, be comprised of similar fatty acids, but migrate further up a TLC plate. Indeed, although rare in mammals, such ether-soluble neutral lipids (with or without alkyl chains), are stored in lipid droplets (Wölk and Fedorova 2024) and migrate to a similar location on TLC, as in our study (Magnusson and Haraldsson 2011; Hutchins, Barkley, and Murphy 2008; Yamazaki et al. 1981). We therefore undertook an extended LC-MS/MS of the UB, in order to describe in detail the types of lipids found at this position.

**Figure 5.**
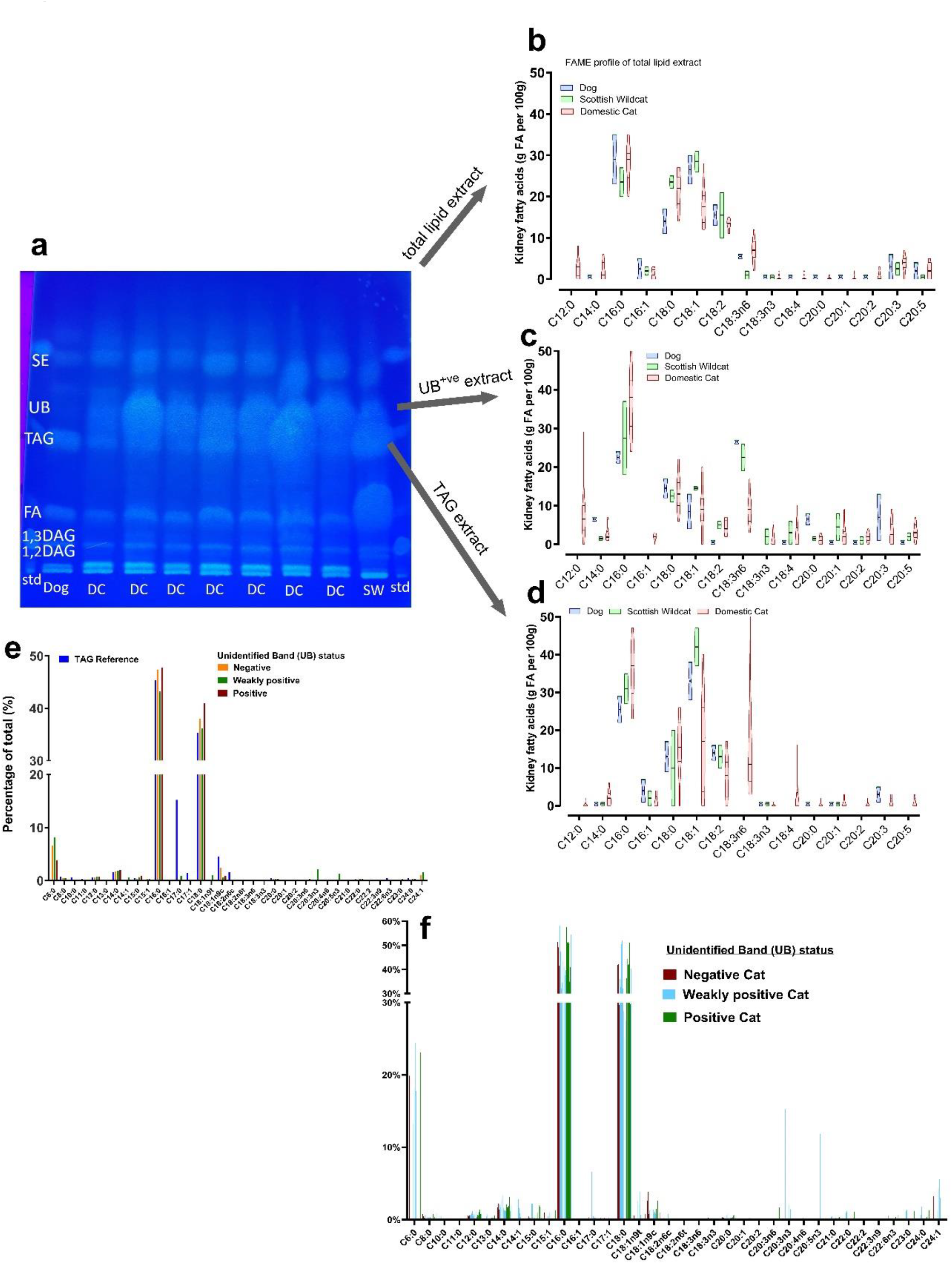
Fatty acid methyl esters (FAME) in domestic cat, dog and Scottish Wildcat kidney extracts. **(a-d) a)** Example TLC plate of separate lipid classes (DC, domestic cat; SW, Scottish Wildcat; FA, fatty acid band, TAG, tryacylglycerides; UB, unidentified band; SE, sterol esters). Each band was scraped and FAME determined from **b)** total lipid extract, **c)** UB^+ve^ band and **d)** TAG band. **e)** data are mean percent of total for FAME from further samples categorised specifically by UB status (UB^-ve^, UB^+ve^ and weakly UB^+ve^),**f)** data are individual values for percent of each FAME for n=18 domestic cats only, with all individuals separated by UB status.

### LC-MS/MS evaluation of the unidentified band

Following extraction and purification of the unidentified band and LC-MS/MS analysis, individuals were categorised according to their TLC result: with (UB^+ve^) or without (UB^-ve^) the UB. Fatty acids were the predominant lipid class in both groups (Figure 6A,B), but UB^+ve^ samples had much greater content of triacylglycerols (‘TG’) and wax esters (WE, Figure 6B). UB^-ve^ samples consistently had a greater proportion of phospholipids, particularly PG (Figure 6A). Expansion of the lipid classes into subclasses using lipidsearch™, then the composition of the UB^-ve^ samples was relatively uniform; predominantly straight chain fatty acids (average: 66.43% [62.84-75.74%], Figure 6C), with relatively greater alkyl-acyl moieties (at ∼6.00%) and fatty ethers (2.18%). In the UB^+ve^ samples these same classes were only 44.65% of total (straight-chain FA), 1.70% (alkyl-acyl lipids), 1.16% (fatty ethers), respectively. In addition, the majority of UB^+ve^ samples had greater content of wax monoesters, unsaturated fatty acids and triacylglycerols, as annoted by lipidsearch™. Interestingly, of note, ‘monoalkyl-diacyl’glycerols were noted in all UB^+ve^, but not UB^-ve^ samples, albeit at low levels Figure 6C). Furthermore, LC-MS revealed that UB^+ve^ extracts were enriched in lipids with atypical compositions and mass profiles, including [O-31:2_23:2], [O-38:7] and [O-4:0_22:6], the ‘O-’ prefix indicates the presence of an ether linkage (Fahy et al. 2007). Such ether-soluble, neutral lipids, particularly those with higher mass values that were present in all UB^+ve^ sample had similar m/z ratio and molecular formula (see Table S1), as the mono-alkyl diacylglycerols (MADAGs) directly identified by (Bartz et al. 2007). Thus, DC samples that were UB^+ve^, in contrast to UB^-ve^ samples consistently had structurally distinct, non-traditional TAG analogues, potentially incorporating ether or branched-chain moieties. We then sought to confirm *in situ*, the presence of such unusual lipids in an unbiased, yet direct, analysis of frozen felid kidney sections with known tubular lipidosis.

**Figure 6.**
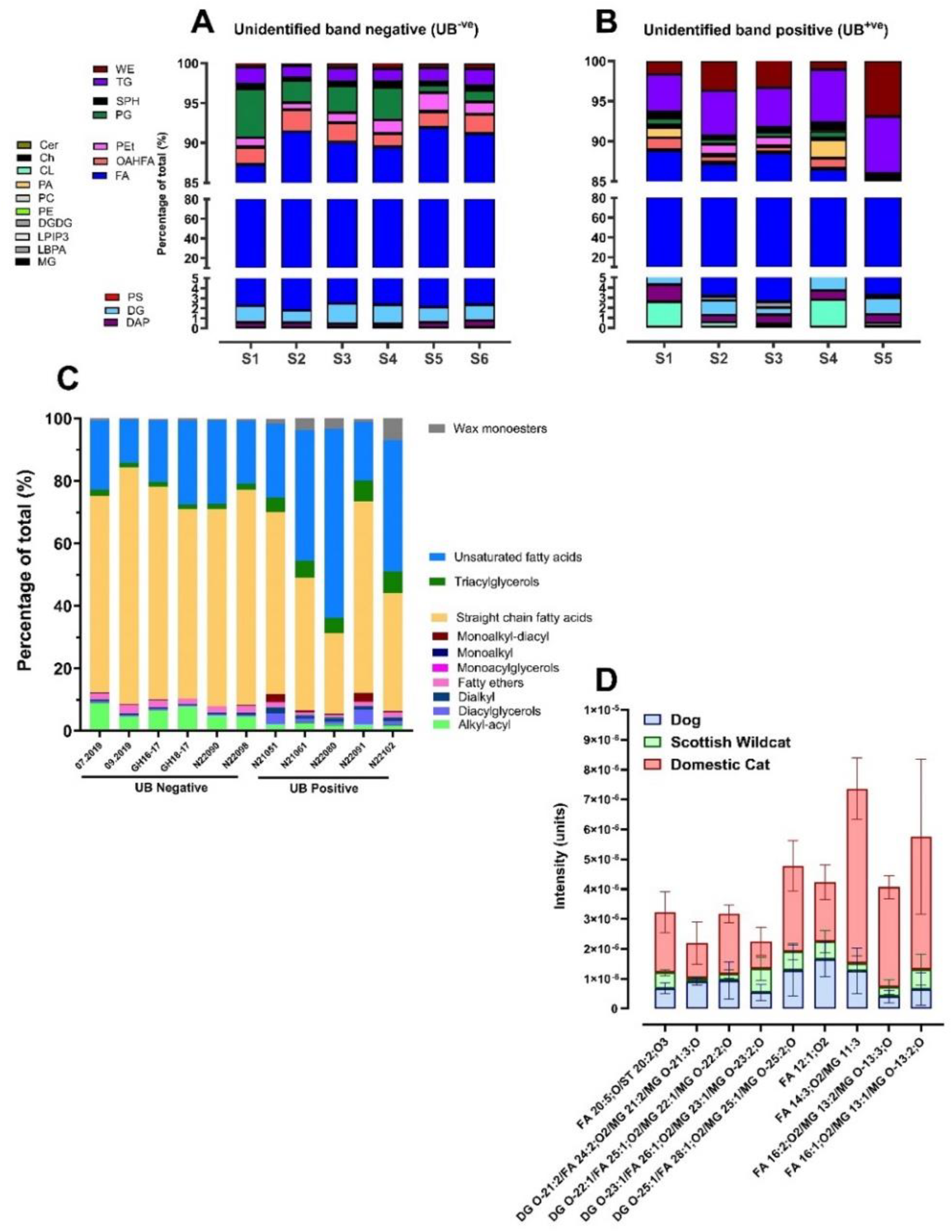
UB lipid composition domestic dog, cat and Scottish Wildcat, as determined by LC-MS and lipid presence in frozen kidney sections by cryo-orbiSIMS. Lipid composition of unidentified band positive versus negative (UB^+ve^, UB^-ve^) in Domestic cat and Scottish Wildcat kidney extracts or frozen sections Lipids determined by LC-MS or cryo-orbiSIMS. Relative percentages of lipid classes present in the unidentified band (UB) across **(A)** n=6 UB^-ve^ and **(B)** n=5 UB^+ve^ individuals, as determined by HPTLC. Cer: ceramide, Ch: cholesterol, CL: cardiolipin, DAP: diacylglycerol pyrophosphate, DG :diacylglycerol, DGDG: digalatosyldiacylglycerol, FA: fatty acid, LPIP3: lysophosphatidylinositol 3-phosphate, LBPA: lysobisphosphatidic acid, MG: monoacylglycerols, OAHFA: (O-acyl)-1-hydroxy fatty acid, PA: phosphatidic acid, PC: phosphatidylcholine, PE: phosphatidylethanolamine, PEt: phosphatidylethanol, PG: phosphatidylglycerol, PS: phosphatidylserine, SPH: sphingosine, TG: triacylglycerol, WE: wax ester. **(C)** Individuals from (A), with percentages of lipid subclass instead, highlighting the distribution of fatty acids into unsaturated fatty acids and straight chain fatty acids. **(D)** Distribution of lipids in frozen kidney from n=4 dog, Scottish wildcat and domestic cat, as determined by cryo-orbiSIMS.

Cryo-OrbiSIMS yielded a spectra of lipids overexpressed in frozen sections of renal cortex (one section per individual: n=4 different samples of dog, DC and SW) from felids verses dog (Figure 6D, Figure S2-5, Table S2). From the n = 2212 peaks detected, OrbiSIMS detected 733 peaks to be unique to domestic cats (cf. Scottish Wildcat and dog) and of those peaks, LipidMaps™ returned 347 lipid assignments (see Supplementary item 9). Many lipids had multiple identifications, having the same elemental composition, e.g. mass: 757.539 (C_42_H_78_O_9_P-) can be identified as PA 39:2;O or PG O-36:3; in this study, the initial annotation is presented. Of note, lipid classes directly identified to be over-expressed in domestic cat, amongst others, included ‘branched-chain fatty acids’, mono-alkylglycerols, and ‘1-alkyl,2-acylglycerophosphoglycerols’ i.e. MADAGS (Figure 6D, Figure S2-5, Table S2).

## Discussion

This paper describes for the first time a comprehensive analysis of kidney tissue lipid content commonly found in companion felids, with dog as a referent category species for the domestic environment, and Scottish Wildcat as a non-domestic referent species. Using Oil-Red-O histochemistry, HPTLC, UPLC-MS/MS, GC-FAME and cryo-OrbiSIMS we have identified and describe a relatively low abundance, yet common for domestic cat (DC), unique profile of renal lipids. Felids in general have abundant Oil-Red-O positive (ORO^+ve^) neutral lipids encapsulated in intracytoplasmic lipid droplets of varying size, usually limited to renal proximal tubule epithelial cells, as previously described (Quimby et al. 2022). Further analysis of these lipid droplets described sub-cellular components full of neutral triacylglycerides (∼72% of total in felids, 39% in dogs), phospholipids (∼17% of total in felids, 32% in dogs) and cholesterol, as expected (Bargmann et al. 1977). However, we additionally report for the first time in a companion animal species, that DC commonly eluted a reproducible, lower-polarity (cf. TAGs) lipid band that was only occasionally present (albeit weakly) in Scottish wildcats, and was absent from dogs and most captive wildcats. The TAG-to-phospholipid ratio in domestic cat, as determined here, is consistent with presence of larger intracytoplasmic lipid droplets formed with a monolayer of phospholipids (mostly phosphatidylcholine). Further lipidomics (LC-MS and spatial cryo-orbiSIMS) and FAME analysis of DC lipid droplets and the unusual lipid band revealed presence of a panoply of modified TAGs, such as odd-and short-chain fatty-acids, branched-chain fatty acids and monoalkyldiacylphospholipids (MADAGS). All modified TAGs would be less polar, by virtue of an ether-linkage, that would account for the unusual banding on lipid chromatography. These data are highly unusual for mammals, particularly when one considers that DC show appearance of renal lipid droplets from an early age. Taken together, the data suggest DC kidney commonly forms unusually modified (ether-linked and/or branched) neutral lipids (such as MADAGs or similar). Definitive structural analysis using reverse-phase HPLC together with NMR would confirm identities. Regardless, given the known association of ectopic lipid including lipid droplets with organ fibrosis, and the known propensity for renal fibrosis in DC (Lawson et al. 2015), then we propose that the propensity for felids to develop tubular lipuria is one of the first pathophysiological outcomes to connect felid biology, diet, age and vulnerability to chronic renal disease.

Evidenced by Oil Red-O and HPTLC, this study confirms that most felids accumulate lipid in their renal cortices, mainly renal proximal tubule epithelial cells (RPTEC) from an early age (Bargmann et al. 1977; Mottram 1916; Modell 1933; Foote and Grafflin 1938; Lobban 1955). Expanding on these studies in cat, we can also report that the lipid droplet lipidome in cat is similar in many respects to experimental animal species; that is, comprised of triacylglycerides, phospholipids and cholesterol (Wölk and Fedorova 2024). A full lipid profile of the cat’s renal intracellular lipid droplets has not been completed previously. Oil Red O was used to stain neutral lipids, including triacylglycerides and diacylglycerides (Koopman, Schaart, and Hesselink 2001), as well as cholesterol esters (Kruth 1984). Analysis of total lipid in felid kidneys indicated that Scottish wildcats, like domestic cats, had evident lipid with a highly variable profile within their kidneys, despite less prevalent histopathologically-determined lipid droplets. Thus, presence of lipid in the kidney cortex appears common amongst felids. However, gradual bioaccumulation with age does not appear to be the case (see Figure 1), consistent with felids regularly exocytosing such droplets into tubular fluid (urine), perhaps explaining the common clinical pathological finding of lipuria in felids, whether domestic (Schwarz et al. 2021) or wild (Hewer, Matthews, and Malkin 1949). Regardless, one feature of the lipids in domestic cat kidneys that is unique, is consistent presence of unusual lipid classes in the droplets, rarely observed in other mammals (Yamazaki et al. 1981), but commonly observed in certain molluscs or squid (Rybin et al. 2017) or in certain conditions of metabolic (i.e. hepatic) stress leading to significant lipolysis (Rico et al. 2021).

### Felid kidneys have an unusually common presence of rarely observed lipids

A distinct lipid band identified by HPTLC as having lower polarity than triacylglycerol and consistently observed in total lipid extracts from domestic cats (referred to as the unidentified band) was an unusual and unexpected finding. The band was absent from extracts of dog kidney, only faintly present in some Scottish wildcats, and appeared unrelated to disease status; kittens, young adults, healthy adult felids with no renal disease or those with diagnosed CKD all generally presented with an UB. Previous studies in other experimental species or under certain conditions had reported an identical band as being specifically, mono-alkyl diacylglycerols (MADAGs) (Bartz et al. 2007; Yamazaki et al. 1981); that is, a TAG where a long-chain fatty alkyl group is connected to the *sn*-1 carbon of glycerol via an ether bond (Magnusson and Haraldsson 2011). MADAGs are essentially storage lipids and ether-analogues of triacylglycerols, and previously have only been noted in any quantity in cells within lipid droplets (Wölk and Fedorova 2024) or where inborn errors of metabolism lead to lysosomal (e.g. Wolman’s) disease (Ma et al. 2017). Quite why felids, particularly domestic cats may be prone to accumulating MADAGs in lipid droplets in the kidney, remains unknown and is beyond the scope of this study, but this information now opens up new avenues of investigation. Are the droplets formed *de novo* with MADAGs coming from diet? Ether lipids are more abundant in squid or oily fish species (Ermolenko et al. 2016) – the by-products of which are often used to flavour domestic cat feeds. However, given the prevalence of MADAGs in the majority of DC in the present study, the fact that they are only observed in tissues with extensive peroxisomes and lysosomes (i.e. they are absent from adipose tissue), suggests formation is local and, in part, due to some aspect of the local cellular microenvironment. Genetic analysis of domestic cat suggested evolutionary adaptation to a diet high in protein and fat – many over-expressed genes were associated with lipid handling (Montague et al. 2014). As ether lipids are either formed and/or broken down by specialised peroxisomal enzymes and other lipases, involving mitochondria and endoplasmic reticulum, and kidneys are highly metabolic organs (similar to heart), suggest that some aspect of felid kidney metabolism lends itself to inappropriate formation of intracellular lipid droplets (Mitrofanova, Merscher, and Fornoni 2023).

The abundant lipids in DC were also enriched with short, mono- and polyunsaturated TAGs such as TG(14:1_14:1_16:1), many of which were completely absent or only minimally present in dogs – which had TAGs mostly enriched with palmitic (16:0), oleic (18:1) and linoleic (18:2) acids (e.g. TG(16:0_16:1_18:1) and TG(16:0_16:0_18:2). Again, such an unusual profile of fatty acids has not been reported before in DC. The profile maybe feline-specific, reflecting some peculiarity of fatty acid synthesis and turnover in the felid kidney, or diet – reflecting intake of a wide variety of different fats or again, some aspect of felid kidney metabolism. Notably, consistent with previous findings in feral cats (Backus, Thomas, and Fritsche 2013), the profile of fatty acids in Scottish wildcat was also much broader than that of dogs, but primarily of longer-chain FA combinations compared to domestic cat (e.g. 16:0, 18:1, 18:2n-6 and 18:3n-3).

In addition, unidentified band positive samples consistently had higher proportions of wax monoesters, and further sub-division elicited a transition from primarily straight chain, saturated fatty acids in UB^-ve^ extracts, to unsaturated fatty acids in UB^+ve^. This suggests a compositional shift in neutral lipid content; specifically, UB^+ve^ samples had a higher proportion of ether-linked lipids such as [O-31:2_23:2], [O-38:7] and [O-4:0_22:6], as well as fatty acid C18:1 (oleic acid). Further confirmation that UB^+ve^ samples tend to be comprised of ether-lipids comes from derivation of the fatty acids using methylation i.e. analysis of fatty-acid methyl esters (FAMEs) data: the protocol for preparation of FAMEs involves cleaving ester, but not ether bonds, UB^+ve^ and UB^-ve^ extracts both had broadly similar percent C16:0 and C18:0 (of total FAMEs). Odd-chain fatty acids (C15:0 and C17:0) were marginally increased in UB^+ve^ samples. This suggests that greater differences in fatty acids between felids with or without an UB is likely to involve non-esterified lipids. This hypothesis is further supported by slightly elevated levels of odd-chain saturated fatty acids, including tridecanoic acid (C13:0) and pentadecanoic acid (C15:0) in UB^+ve^ individuals. Presence of these odd-chain lipids in felid kidneys could indicate higher intake of branched-chain amino- or fatty acids (BCFAs) in those felids that are UB^+ve^ or, as is more likely, *de novo* accumulation. The latter can occur when cells (usually liver) become deplete in certain micronutrients or excessive in others such as the short-chain fatty acids (proprionate, valerate). For example, in ovine white liver disease (a condition affecting sheep caused by cobalt deficiency, leading to vitamin B_12_ deficiency), the liver accumulates increased levels of branched-chain fatty acids, particularly propionate, due to disruptions in the metabolic pathway that utilizes these fatty acids, resulting in a characteristic pale, fatty liver (Kennedy et al. 1994). The extent to which felids may exhibit a similar cellular microenvironment, from an early age, is not known.

Here, direct, unbiased and non-destructive analysis of renal cortical tissue using cryo-orbiSIMS provided further support for a species-specific lipid signature in kidneys. Domestic cat kidneys had 733 unique peaks, compared to dog and wildcat, 347 of which were assigned lipid identities. Among these were elevated levels of fatty acids, diacylglycerols, and monoacylglycerols – many of which were ether-linked (O-, see Supplementary item 4). Indeed, n=19 had accurate masses previously, positively identified as MADAGS (Bartz et al. 2007). The presence of differing lipids classes in lipid droplets of the same feline kidney tissue further suggests heterogeneity in lipid storage, potentially reflecting cell-specific lipid metabolism or sequestration, not surprisingly given the heterogeneity of cell types in kidney cortex. The distinct spatial distribution, combined with the detection of ether-linked phospholipids reinforces the idea that feline kidneys may preferentially accumulate atypical lipid structures as either an adaptive or pathological response to chronic metabolic stress.

### Age and sex of the animal

Age was an important consideration in this study as a more lipidic phenotype emerged from kitten through adult to senior cats, as previously observed (Quimby et al. 2022). While the phenotype of tubular lipidosis is often macroscopically indistinct (Lobban 1955), the propensity to form lipid droplets appears to increase with age – 100 % of cats ≥15 yrs had tubular lipidosis, according to (Quimby et al. 2022). We observed the overall fatty acid profile of young and senior cat kidneys to be similar, but the propensity for TAG to migrate and become UB^+ve^ was notable; young cats generally had a prominent TAG band with sometimes a weak UB^+ve^ or ‘MADAG’ band. In contrast, for senior cats then presence of a prominent UB^+ve^ or ‘MADAG’ band tended to be associated with a weak TAG band; our interpretation being that with seniority, the renal cells become more predisposed to the conditions that exacerbate formation of intracellular lipid droplets with a unique lipid signature, as previously described here. The two oldest cats in this study were 13-years and 15-years of age females, both were UB^+ve^. In contrast, a male Scottish wildcat kitten and a 7-month-old female were both UB^-ve^. Regardless, it is accepted across animal models, experimental or human, that deposition of ectopic lipid (i.e. lipid not stored in adipose tissue) is toxic to cells, including renal (Izquierdo-Lahuerta, Martínez-García, and Medina-Gómez 2016; Weinberg 2006). The relatively high lipid content of the domestic cat kidney, evidenced as early as young adulthood, whereas first stage symptoms of renal disease do not become evident until approximately 7-8 years of age, would suggest a role for tubular lipidosis in the aetiopathogenesis of chronic kidney disease (CKD) in felids, a hitherto unforeseen and overlooked cause as yet.

In summary, felids fundamentally appear prone to forming intracellular lipid droplets in their metabolically active renal cells from an early age. The lipid droplets have a unique signature in domestic cat, distinguished by commonly having, what otherwise would be rare in mammals, the presence of unique fatty acids (e.g. odd-chain, branched) and lipids (e.g. MADAGS) that likely reflect the local cellular microenvironment. Clearly, with lipiduria commonly referred to as ‘incidental’ in domestic cat, then further work should determine if the lipids in feline urine are the same as those found in lipid droplets in kidney cells (i.e. reflecting regular turnover). Regardless, ectopic lipid is always deleterious. We suggest that tubular lipidosis is one of the first indicators of a renal cellular environment that, years later, is manifest as CKD.

## Acknowledgements and Financial Disclosure

The authors would like to acknowledge Georg Hantke and Andrew Kitchener (National Museums Scotland) for providing samples of Scottish and captive Wildcat kidneys; multiple anatomic pathologists in The School of Veterinary Medicine & Science pathology department for assistance with domestic feline and canine samples; Judith Ngere (Thermo Fisher Scientific) for help with Lipidsearch™; Drs Richard Broughton and Louise Michaelson (Rothampsted Research Station) for initial TLC, FAME & LC-MS/MS analyses; Drs Vincenzo Di Bari and Ruth Price for assistance with HPTLC analysis and interpretation, Drs Louise Williams and Noriane Cochetel for GC-MS analysis, Dongfang Li for FAME analysis. Professors Dong-Hyun Kim and David Scurr for help with OrbiSIMS and mass spectrometry. This study was funded in part by Dechra Veterinary Products through a studentship to R. Alborough (2017 – 2020) and in part by the Biotechnology and Biological Sciences Research Council (BBSRC) as part of the University of Nottingham DTP PhD studentship awarded to R.A Brociek (Grant code: RS86P5).

## References

Backus, Robert C, David G Thomas, and Kevin L Fritsche. 2013. ‘Comparison of inferred fractions of n-3 and n-6 polyunsaturated fatty acids in feral domestic cat diets with those in commercial feline extruded diets‘, Am J Vet Res, 74: 589–97.

Bargmann, W., B. Krisch, H. Leonhardt, and M. Mályusz. 1977. ‘Lipids in the proximal convoluted tubule of the cat kidney and the reabsorption of cholesterol‘, Cell and Tissue Research, 177: 523–38.

Bartz, René, Wen-Hong Li, Barney Venables, John K Zehmer, Mary R Roth, Ruth Welti, Richard GW Anderson, Pingsheng Liu, and Kent D Chapman. 2007. ‘Lipidomics reveals that adiposomes store ether lipids and mediate phospholipid traffic1, s?‘, Journal of Lipid Research, 48: 837–47.

Bobulescu, Ion Alexandru. 2010a. ‘Renal lipid metabolism and lipotoxicity‘, Current opinion in nephrology and hypertension, 19: 393–402.

Bobulescu, Ion Alexandru. 2010b. ‘Renal lipid metabolism and lipotoxicity‘, Current opinion in nephrology and hypertension, 19: 393.

Bostrom, P., L. Andersson, M. Rutberg, J. Perman, U. Lidberg, B. R. Johansson, J. Fernandez-Rodriguez,J. Ericson, T. Nilsson, J. Boren, and S. O. Olofsson. 2007. ’sNARE proteins mediate fusion between cytosolic lipid droplets and are implicated in insulin sensitivity‘, Nat Cell Biol, 9: 1286–93.

Chapman, Kent D., John M. Dyer, and Robert T. Mullen. 2012. ‘Biogenesis and functions of lipid droplets in plants: Thematic Review Series: Lipid Droplet Synthesis and Metabolism: from Yeast to Man‘, Journal of Lipid Research, 53: 215–26.

D‘Arcy, Rachel. 2018. ‘Chronic kidney disease in non-domestic felids in Australian zoos‘.

Ducharme, N. A., and P. E. Bickel. 2008. ‘Lipid droplets in lipogenesis and lipolysis‘, Endocrinology, 149: 942–9.

Else, P. L., and A. J. Hulbert. 1985. ‘An allometric comparison of the mitochondria of mammalian and reptilian tissues: the implications for the evolution of endothermy‘, J Comp Physiol B, 156: 3–11.

Ermolenko, Ekaterina, Nikolay Latyshev, Ruslan Sultanov, and Sergey Kasyanov. 2016. ‘Technological approach of 1-O-alkyl-sn-glycerols separation from Berryteuthis magister squid liver oil‘, Journalof food science and technology, 53: 1722–26.

Escasany, Elia, Adriana Izquierdo-Lahuerta, and Gema Medina-Gómez. 2018. ‘Chapter 7 - Kidney Damage in Obese Subjects: Oxidative Stress and Inflammation.‘ in Amelia Marti del Moral and Concepción María Aguilera García (eds.), Obesity (Academic Press).

Fahy, Eoin, Manish Sud, Dawn Cotter, and Shankar Subramaniam. 2007. ‘LIPID MAPS online tools for lipid research‘, Nucleic Acids Research, 35: W606–W12.

Foote, John J, and Allan L Grafflin. 1938. ‘Quantitative measurements of the fat-laden and fat-free segments of the proximal tubule in the nephron of the cat and dog‘, The Anatomical Record, 72: 169–79.

Hewer, TF, L Harrison Matthews, and T Malkin. 1949. “Lipuria in tigers.” In Proceedings of the Zoological Society of London, 924–28. Wiley Online Library.

Hutchins, Patrick M, Robert M Barkley, and Robert C Murphy. 2008. ’separation of cellular nonpolar neutral lipids by normal-phase chromatography and analysis by electrospray ionization mass spectrometry‘, Journal of Lipid Research, 49: 804–13.

Izquierdo-Lahuerta, Adriana, Cristina Martínez-García, and Gema Medina-Gómez. 2016. ‘Lipotoxicity as a trigger factor of renal disease‘, Journal of nephrology, 29: 603–10.

Kang, Hyun Mi, Seon Ho Ahn, Peter Choi, Yi-An Ko, Seung Hyeok Han, Frank Chinga, Ae Seo Deok Park, Jianling Tao, Kumar Sharma, James Pullman, Erwin P. Bottinger, Ira J. Goldberg, and Katalin Susztak. 2014. ‘Defective fatty acid oxidation in renal tubular epithelial cells has a key role in kidney fibrosis development‘, Nature Medicine, 21: 37.

Kennedy, DG, S Kennedy, WJ Blanchflower, JM Scott, DG Weir, AM Molloy, and PB Young. 1994. ‘Cobalt-vitamin B12 deficiency causes accumulation of odd-numbered, branched-chain fatty acids in the tissues of sheep‘, British Journal of Nutrition, 71: 67–76.

Koopman, René, Gert Schaart, and Matthijs K Hesselink. 2001. ‘Optimisation of oil red O staining permits combination with immunofluorescence and automated quantification of lipids‘, Histochemistry and cell biology, 116: 63–68.

Kotowska, Anna M., Junting Zhang, Alessandro Carabelli, Julie Watts, Jonathan W. Aylott, Ian S. Gilmore, Paul Williams, David J. Scurr, and Morgan R. Alexander. 2023. ‘Toward Comprehensive Analysis of the 3D Chemistry of Pseudomonas aeruginosa Biofilms‘, Analytical Chemistry, 95: 18287–94.

Kruth, HS. 1984. ‘Localization of unesterified cholesterol in human atherosclerotic lesions. Demonstration of filipin-positive, oil-red-O-negative particles‘, The American Journal of Pathology, 114: 201.

Lawson, Jack, Jonathan Elliott, Caroline Wheeler-Jones, Harriet Syme, and Rosanne Jepson. 2015. ‘Renal fibrosis in feline chronic kidney disease: Known mediators and mechanisms of injury‘, The Veterinary Journal, 203: 18–26.

Lobban, Mary C. 1955. ’some observations on the intracellular lipid in the kidney of the cat‘, Journal of Anatomy, 89: 92.

Lucke, Vanda M. 1968. ‘Renal disease in the domestic cat‘, The Journal of Pathology and Bacteriology, 95: 67–91.

Ma, Zhengping, Joelle M. Onorato, Luping Chen, David W. Nelson, Chi-Liang Eric Yen, and Dong Cheng. 2017. ’synthesis of neutral ether lipid monoalkyl-diacylglycerol by lipid acyltransferases[S]‘, Journal of Lipid Research, 58: 1091–99.

Macnider, Wm Deb. 1945. ‘Occurrence of Stainable Lipoid Material in Renal Epithelium of Animals Falling in Different Age Segments‘, Proceedings of the Society for Experimental Biology and Medicine, 58: 326–28.

Magnusson, C. D., and G. G. Haraldsson. 2011. ‘Ether lipids‘, Chem Phys Lipids, 164: 315–40.

Martino-Costa, A. L., F. Malhão, C. Lopes, and P. Dias-Pereira. 2017. ‘Renal Interstitial Lipid Accumulation in Cats with Chronic Kidney Disease‘, J Comp Pathol, 157: 75–79.

Maunsbach, Arvid B., and Claes Wirsén. 1966. ‘Ultrastructural changes in kidney, myocardium and skeletal muscle of the dog during excessive mobilization of free fatty acids‘, Journal of Ultrastructure Research, 16: 35–54.

Mitrofanova, Alla, Sandra Merscher, and Alessia Fornoni. 2023. ‘Kidney lipid dysmetabolism and lipid droplet accumulation in chronic kidney disease‘, Nature Reviews Nephrology, 19: 629–45.

Modell, Walter. 1933. ‘Observations on the lipoids in the renal tubule of the cat‘, The Anatomical Record, 57: 13–27.

Montague, Michael J., Gang Li, Barbara Gandolfi, Razib Khan, Bronwen L. Aken, Steven M. J. Searle, Patrick Minx, LaDeana W. Hillier, Daniel C. Koboldt, Brian W. Davis, Carlos A. Driscoll, Christina S. Barr, Kevin Blackistone, Javier Quilez, Belen Lorente-Galdos, Tomas Marques-Bonet, Can Alkan, Gregg W. C. Thomas, Matthew W. Hahn, Marilyn Menotti-Raymond, Stephen J. O’Brien, Richard K. Wilson, Leslie A. Lyons, William J. Murphy, and Wesley C. Warren. 2014. ‘Comparative analysis of the domestic cat genome reveals genetic signatures underlying feline biology and domestication‘, Proceedings of the National Academy of Sciences, 111: 17230–35.

Mottram, VH. 1916. ‘Fatty infiltration of the cat’s kidney‘, The Journal of Physiology, 50: 380.

Newkirk, K. M., S. J. Newman, L. A. White, B. W. Rohrbach, and E. C. Ramsay. 2011. ‘Renal Lesions of Nondomestic Felids‘, Vet Pathol, 48: 698–705.

Olofsson, Sven-Olof, Pontus Boström, Linda Andersson, Mikael Rutberg, Jeanna Perman, and Jan Borén. 2009. ‘Lipid droplets as dynamic organelles connecting storage and efflux of lipids‘, Biochimica et Biophysica Acta (BBA) - Molecular and Cell Biology of Lipids, 1791: 448–58.

Pan, D. A., A. J. Hulbert, and L. H. Storlien. 1994. ‘Dietary fats, membrane phospholipids and obesity‘, J Nutr, 124: 1555–65.

Quimby, Jessica M, Shannon M McLeland, Rachel E Cianciolo, Katharine F Lunn, Jody P Lulich, Andrea Erickson, and Lara B Barron. 2022. ‘Frequency of histologic lesions in the kidneys of cats without kidney disease‘, J Feline Med Surg, 24: e472–e80.

Rico, Jorge Eduardo, Sina Saed Samii, Yu Zang, Pragney Deme, Norman J. Haughey, Ester Grilli, and Joseph W. McFadden. 2021. ‘Characterization of the Plasma Lipidome in Dairy Cattle Transitioning from Gestation to Lactation: Identifying Novel Biomarkers of Metabolic Impairment‘, Metabolites, 11: 290.

Rybin, Viacheslav G, Andrey B Imbs, Darja A Demidkova, and Ekaterina V Ermolenko. 2017. ‘Identification of molecular species of monoalkyldiacylglycerol from the squid Berryteuthis magister using liquid chromatography–APCI high-resolution mass spectrometry‘, Chem Phys Lipids, 202: 55–61.

Sakuma, Ikki, Rafael C. Gaspar, Ali R. Nasiri, Sylvie Dufour, Mario Kahn, Jie Zheng, Traci E. LaMoia, Mateus T. Guerra, Yuki Taki, Yusuke Kawashima, Dean Yimlamai, Mark Perelis, Daniel F. Vatner, Kitt Falk Petersen, Maximilian Huttasch, Birgit Knebel, Sabine Kahl, Michael Roden, Varman T. Samuel, Tomoaki Tanaka, and Gerald I. Shulman. 2025. ‘Liver lipid droplet cholesterol content is a key determinant of metabolic dysfunction–associated steatohepatitis‘, Proceedings of the National Academy of Sciences, 122: e2502978122.

Schwarz, T., E. Shorten, M. Gennace, J. Saunders, M. Longo, F. S. Costa, M. Parys, and D. Gunn-Moore. 2021. ‘CT features of feline lipiduria and renal cortical lipid deposition‘, J Feline Med Surg, 23: 357–63.

Stannard, S. R., and N. A. Johnson. 2004. ‘Insulin resistance and elevated triglyceride in muscle: more important for survival than ‘thrifty’ genes?’, J Physiol, 554: 595–607.

Starr, Nichola J, Mohammed H Khan, Max K Edney, Gustavo F Trindade, Stefanie Kern, Alexander Pirkl, Matthias Kleine-Boymann, Christopher Elms, Mark M O‘Mahony, and Mike Bell. 2022. ‘Elucidating the molecular landscape of the stratum corneum‘, Proceedings of the National Academy of Sciences, 119: e2114380119.

Stich, V., and M. Berlan. 2004. ‘Physiological regulation of NEFA availability: lipolysis pathway‘, Proceedings of the Nutrition Society, 63: 369–74.

Thannickal, Victor J., Yong Zhou, Amit Gaggar, and Steven R. Duncan. 2014. ‘Fibrosis: ultimate and proximate causes‘, The Journal of Clinical Investigation, 124: 4673–77.

Weinberg, J. M. 2006. ‘Lipotoxicity‘, Kidney International, 70: 1560–66.

Whitby, A., P. Pabla, B. Shastri, L. Amugi, Á Del Río-Álvarez, D. H. Kim, L. Royo, C. Armengol, and M. Dandapani. 2023. ‘Characterisation of Aberrant Metabolic Pathways in Hepatoblastoma Using Liquid Chromatography and Tandem Mass Spectrometry (LC-MS/MS)‘, Cancers (Basel), 15.

Wölk, Michele, and Maria Fedorova. 2024. ‘The lipid droplet lipidome‘, FEBS Letters, 598: 1215–25.

Wynn, Thomas A., and Thirumalai R. Ramalingam. 2012. ‘Mechanisms of fibrosis: therapeutic translation for fibrotic disease‘, Nat Med, 18: 1028–40.

Yamazaki, Takeshi, Yousuke SEYAMA, Hideaki OTSUKA, Hideko OGAWA, and Tamio YAMAKAWA. 1981. ‘Identification of alkyldiacylglycerols containing saturated methyl branched chains in the Harderian gland of guinea pig‘, The Journal of Biochemistry, 89: 683–91.

