## Supplementary file for "Lipid droplets in felid kidneys: prevalence and composition by lipidomics"

### Supplementary Information

1. **Figure S1:** Summary of methods to extract and process lipids from cat kidneys
2. Item 2: expanded legend for Figure 1A, B, C, D
3. **Table S1**: LC-MS analysis of UB from current data in comparison to known MADAGS from previous publications
4. **Figure S2**. Lipids identified as significantly different by cryo-orbiSIMS between Domestic Cat, Scottish Wildcat and Domestic dog.
5. **Figure S3**. Spatial distribution of the lipid peaks statistically significantly more intense in Domestic Cat (cf. Scottish Wildcat, Domestic dog) but not co-localised
6. **Figure S4**. Spatial distribution of the lipid peaks statistically significantly more intense in Domestic Cat (cf. Scottish Wildcat, Domestic dog) and co-localised
7. **Table S2**: Selected lipids unique to domestic cat (cf. domestic dog, Scottish wildcat) as determined by cryoorbiSIMS
8. **Figure S5**: Spatial distribution of top 20 lipids exclusive to domestic cat by cryo-orbiSIMS
9. LipidMaps assignment of 347 peaks unique to domestic cat (see excel file, n=347 lipids exclusive to DC_OrbiSIMS.xlsx)

**Figure S1.** Summary of method to extract and process lipids from cat kidneys

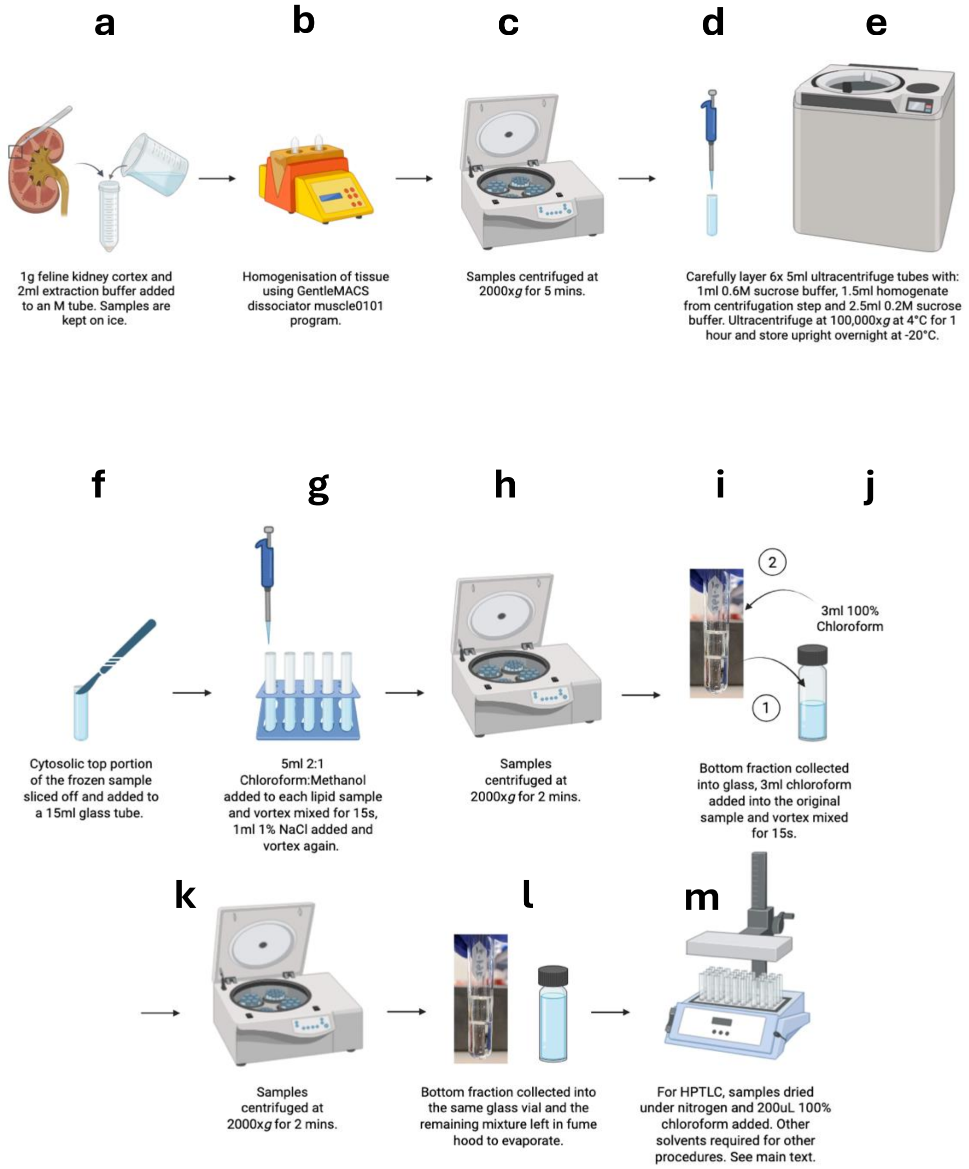


**Figure S1.** Lipid extraction, based on a modified Bligh & Dyer method. **(a)** 1g of frozen renal cortex tissue is mixed with 0.6M protease-free extraction buffer and **(b)** homogenised using the GentleMACS$^{TM}$ dissociator. After, **(c)** homogenate is centrifuged at 2000g for 5 minutes and **(d)** a sucrose step gradient prepared in 5ml polyallomer tube by overlaying 1mL 0.6M protease-free sucrose extraction buffer, 1.5mL of centrifuged homogenate and 2.5mL of 0.25M protease-free sucrose extraction buffer before **(e)** ultracentrifugation at 100,000g at 4°C for 1hr and stored at -20°C overnight. **(f)** The top portion of the frozen extract was transferred to a 15mL glass tube and **(g)** 5mL Cholorform:Methanol (2:1) added to each before vortex mixing for 15s. 1mL NaCl was added and mixed for a further 15s before **(h)** centrifuging at 2000g for 2 minutes. **(i)** The lower organic phase, comprising the sample lipids, was transferred into a new glass vial and **(j)** a further 2 ml of 100% chloroform was added to the original sample, vortexed twice for 15 seconds and centrifuged at **(k)** 1000*g* for 2 minutes. **(l)** The lower layer was transferred and pooled in the new vial and **(m)** the contents were concentrated by drying under nitrogen gas and re-suspending in 200µl 100% chloroform. Samples were stored in vials at -80°C.

### Supplementary Item 2: Expanded legend for Figure 1A, B, C, D

**Figure 1A**. Separation of (left to right) n=6 Scottish wildcats (2-7), n=3 domestic cats (8-10) and n=3 domestic dogs (11-13). Arrow indicates the location of the unknown band. Presence of unidentified band in individuals 2-4, 6, and 8-10. Distinct TAG bands were seen in lanes 5 and 7; a Scottish wildcat adult and kitten, lane 9; 15.5yo domestic cat with a history of CKD and lane 13; a 5.5yo severely overweight Alaskan malamute. 10uL was applied for all samples. Lanes 1 and 14 contain 5uL MAG/DAG/TAG standard.

**Figure 1B**. Separation of (left to right) n=5 domestic cats (2-6), n=4 Scottish wildcats (7-10) and n=2 zoo wildcats (11-12). Arrow indicates the location of the unknown band. Presence of unknown band in individuals 2,3,5 and small amount in 6-9. Distinct TAG bands were seen in lane 6; a 7-month-old domestic cat kitten, and lanes 11 and 12; a snow leopard of unknown age, and an 18yo tiger. 10uL was applied for all samples. Lanes 1 and 13 contain 5uL MAG/DAG/TAG standard.

**Figure 1C** Separation of (left to right) 2) Scottish wildcat adult, 3) Scottish wildcat junior, 4) Scottish wildcat junior, 5) Scottish wildcat adult, 6) Scottish wildcat junior and 7) snow leopard (15 years old). Juniors of unknown age are classified based on body size and are expected to my 1 year or less. Arrow indicates the location of the unknown band. Presence of unknown band in all individuals. More defined TAG bands were seen in lanes 3 and 6; two domestic cat kittens (24 weeks and 1 year, respectively). 10uL was applied for all samples. Lanes 1 and 8 contain 5uL MAG/DAG/TAG standard.

**Figure 1D** Separation of (left to right) 2) domestic cat (adult), 3) domestic cat (kitten-24 weeks), 4) domestic cat (adult), 5) domestic cat (adult), 6) domestic cat (junior-1 year), 7) Scottish wildcat (junior-1 year). Juniors of unknown age are classified based on body size and are expected to my 1 year or less. Arrow indicates the location of the unknown band. Presence of unknown band in all individuals. More defined TAG bands were seen in lanes 3 and 6; two domestic cat kittens (24 weeks and 1 year, respectively). 10uL was applied for all samples. Lanes 1 and 8 contain 5uL MAG/DAG/TAG standard.

**Table S1:** LC-MS analysis of UB from current data in comparison to known MADAGS from previous publications

| Mass:charge ratio (m/z) with of MADAGS from (Bartz et al. 2007) | | OrbiSIMS result (Kidney section, cat) | LC-MS of selected unidentified band (UB) lipids | | |
|---|---|---|---|---|---|
| m/z | Structure | m/z | Calc m/z | Lipid Molecule | Ion |
| 804.7 | 48:3 | 804.4947<br>804.5554 | 804.6800 | TG(47:2)+O:(s) | $M^+NH_4$ |
| 856.8 | 52:5 | 856.4594 | 856.6193 | PG(23:3_19:0) | $M^+NH_4$ |
| 860.8 | 52:3 | 860.5381 | 859.7394 | PC(O-42:0) | $M^+H$ |
| 886.8 | 54:4 | 886.5533<br>886.5707 | 886.6687 | TG(54:10)+O:(s) | $M^+NH_4$ |
| 890.9 | 54:2 | 890.5846 | 890.5098 | PG(46:14) | $M^+Na$ |
| 944.9 | 58:3 | 944.5406 | 944.8044 | TG(56:3)+OO:(s) | $M^+H$ |
| 948.9 | 58:1/58:8 | 948.7398 | 948.5810 | DGDG(37:7) | $M^+Na$ |

**Table S1.** Comparison of accurate lipid masses observed in the current paper (Brociek et al 2025) from cryoOrbiSIMS of n=4 independent cryo-sections (10μm) of domestic cat with known MADAGS positively identified by (Bartz et al. 2007). Calculated m/z is ...

### References


Bartz, René, Wen-Hong Li, Barney Venables, John K Zehmer, Mary R Roth, Ruth Welti, Richard GW Anderson, Pingsheng Liu, and Kent D Chapman. 2007. 'Lipidomics reveals that adiposomes store ether lipids and mediate phospholipid traffic1, s', *Journal of Lipid Research*, 48: 837-47.

**Figure S2**. Lipids identified as significantly different by cryo-orbiSIMS between Domestic Cat, Scottish Wildcat and Domestic dog.

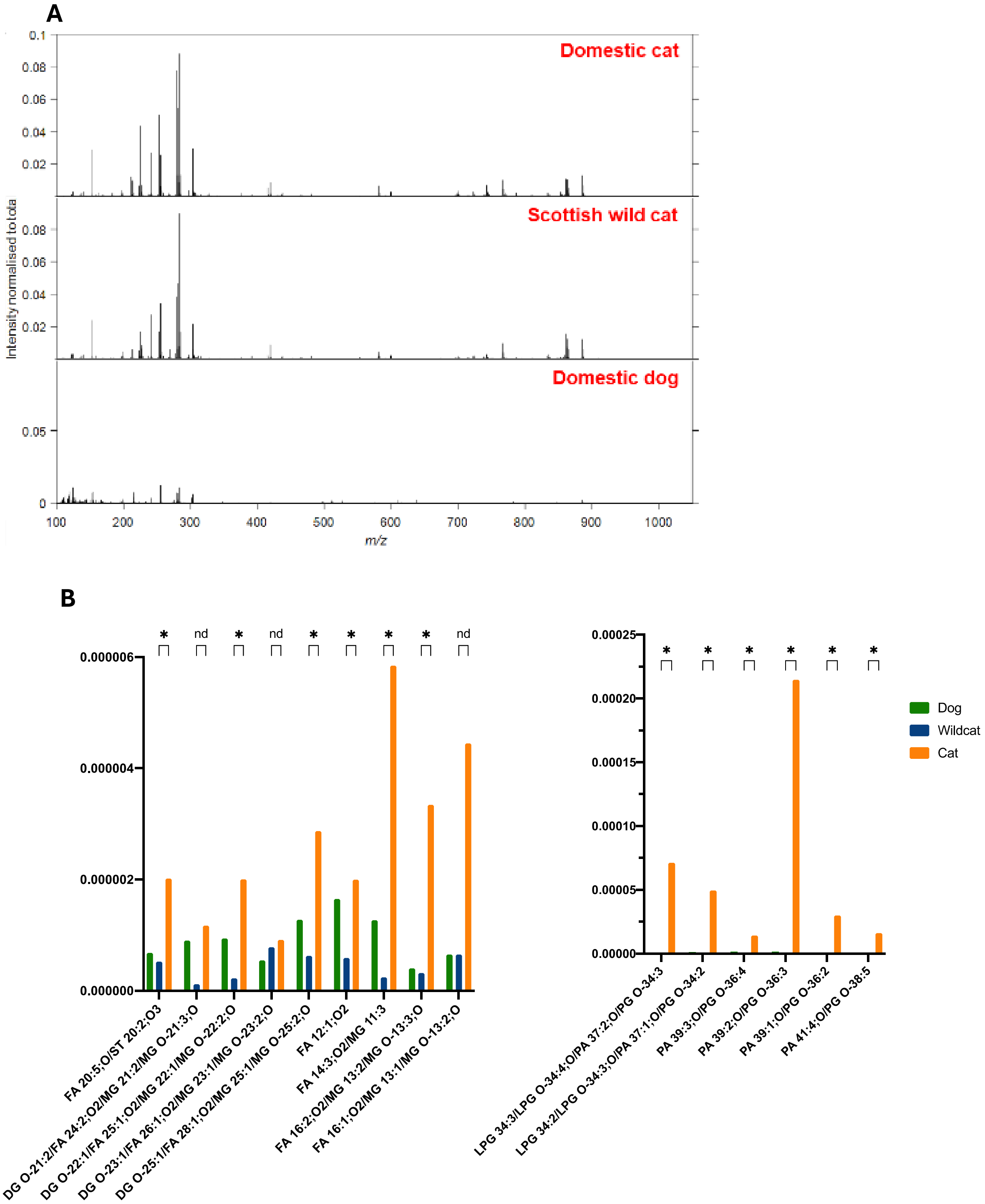


**Figure S2. A)** Assessment of n=4 domestic cat, n=4 dog and n=4 Scottish wildcat kidney spectra show similarity between domestic cats and Scottish wildcats, in comparison to domestic dog – which is characterised by different chemistrys. **B)** demonstrates lipid assignments which were found to be more prevalent in the domestic cat kidney section compared to dog or wildcat kidney sections. Statistical significance based on an unpaired t-test is highlighted between the intensities of the lipids in the wildcat sample versus the domestic cat sample: these were assigned as fatty acids, diglycerols or monoglycerols. Additionally, a group of lipids in the higher mass region was found to be significantly more intense in the domestic cat sample, these belong predominantly to diacylglycerophosphates or diacylglycerophosphoglycerols (these groups of lipids have similar elemental compositions and as such they are often isobaric and cannot be distinguished by the accurate mass alone).

**Figure S3**. Spatial distribution of the lipid peaks statistically significantly more intense in Domestic Cat (cf. Scottish Wildcat, Domestic dog) but not co-localised

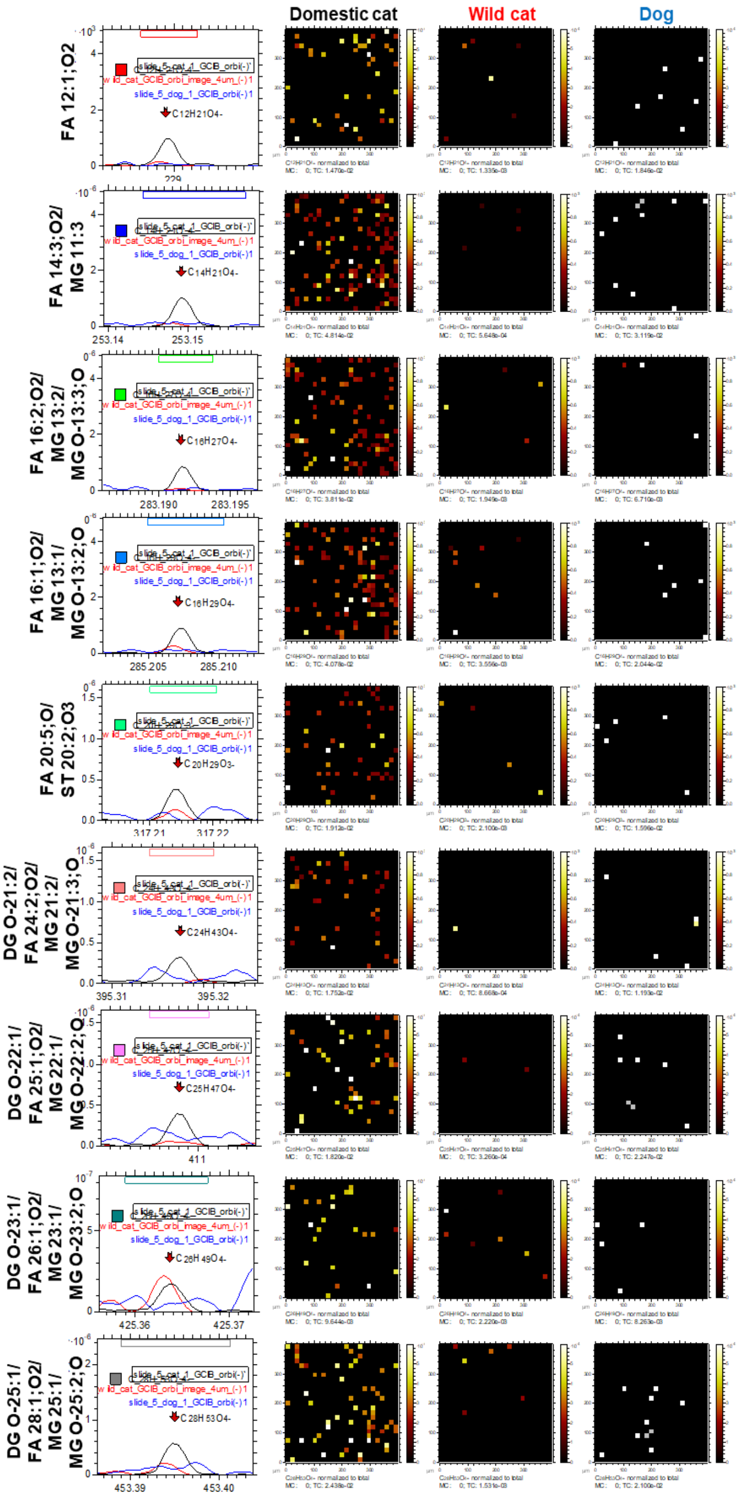


**Figure S3.** These lipid peaks were not generally co-localised within the analysis window of 400 μm × 400 μm and were identified as fatty acids (FA), mono-glycerols (MG), mono-alkyldiacylglycerols (MG O-; ‘MADAGS’) or diacylglycerols (DG).

**Figure S4**. Spatial distribution of the lipid peaks statistically significantly more intense in Domestic Cat (cf. Scottish Wildcat, Domestic dog) that were co-localised

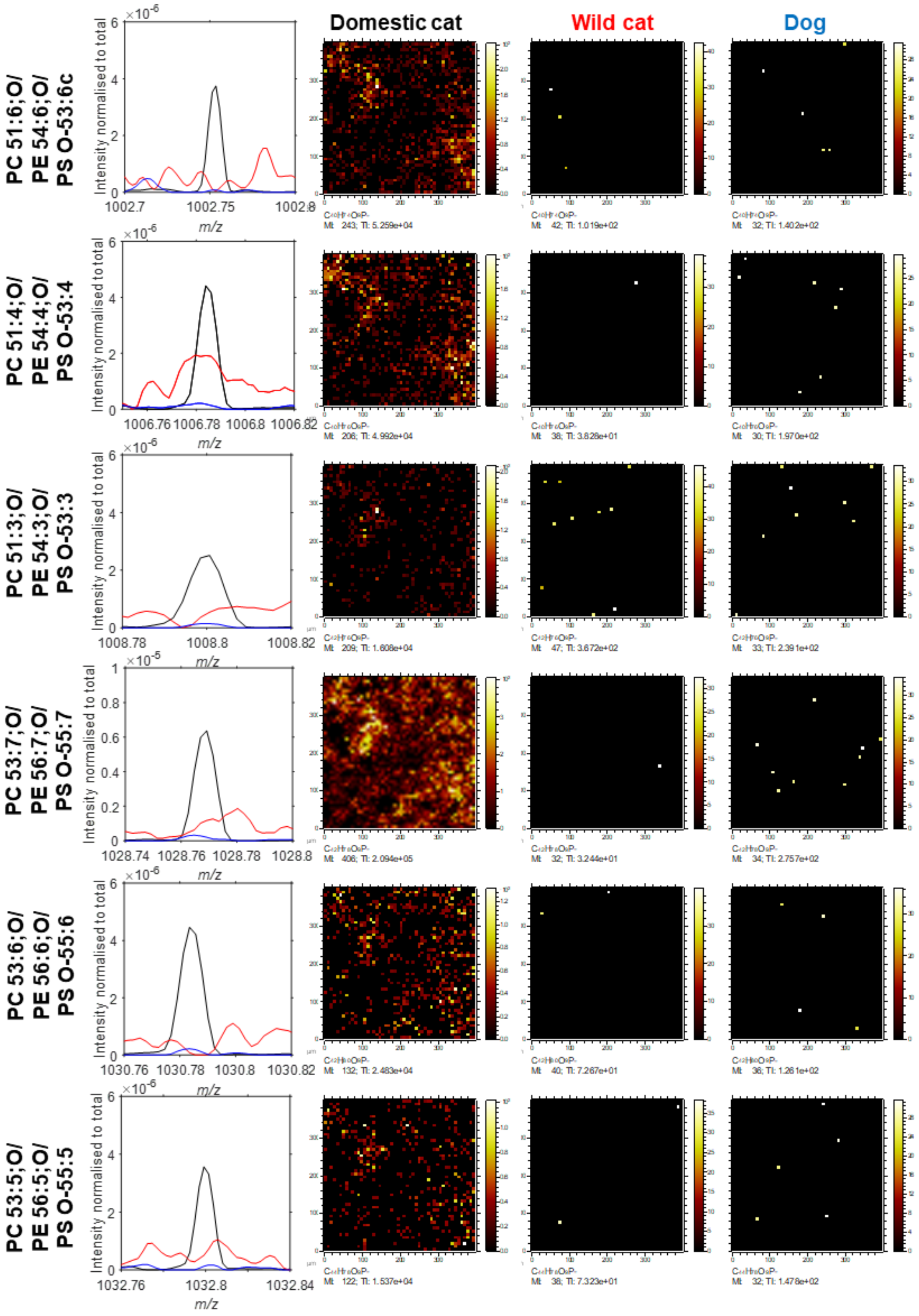


**Figure S4.** Lipid peaks generally co-localised within the analysis window of 400 μm × 400 μm belonging to groups such as diacylglycerophosphocholines (PC), diacylglycerophosphoethanolamines (PE) or diacylglycerophosphoserines (PS).

**Table S2: Lipids unique to domestic cat (cf. domestic dog, Scottish wildcat) as determined by cryoorbiSIMS.**

| Mass (u) | Potential structure | Delta score (±0.001m/z) † | Ion | Structure | Name | Class |
|---|---|---|---|---|---|---|
| 229.1447 | FA 12:1;O2 | .0013 | [M+H]+ | FA 12:2;O2 | 9-hydroxy-traumatin | Hydroxy Fatty Acids |
| 253.1446 | FA 14:3;O2/MG 11:3 | .0036 | [M+Na]+ | FA 12:1;O2 | Dodecanedioic acid | Dicarboxylic Acids |
| 283.1917 | FA 16:2;O2/MG 13:2/MG O-13:3;O | .0037 | [M+Na]+ | FA 14:0;O2 | Ipurolic acid | Hydroxy Fatty Acids |
| 285.2073 | FA 16:1;O2/MG 13:1/MG O-13:2;O | .0140*<br>.0351*<br>.0248* | [M+H]+<br>[M+H]+<br>[M+Na]+ | ST 20:3;O<br>FA 17:1;O<br>FA 17:4 | Lynestrenol<br>16-oxo-heptadecanoic acid<br>14:4(2E,4E,8E,10E)(6Me[S],9Me,12Me[S]) | C21 steroids and derivatives<br>Oxo Fatty Acids<br>Branched Chain Fatty Acids |
| 317.2121 | FA 20:5;O/ST 20:2;O3 | .0010<br>.0010<br>.0034 | [M+H]+<br>[M+H]+<br>[M+Na]+ | FA 20:6;O<br>ST 20:3;O3<br>FA 18:3;O | 15-deoxy-delta-12,14-PGA2<br>2-Methoxyestradiol-3-methylether<br>12-keto-8E,10E-octadecadienoic acid | Eicosanoids- Prostaglandins<br>C18 Steroid (estrogens) and derivatives<br>Unsaturated Fatty Acid |
| 395.3166 | DG O-21:2/FA 24:2;O2/MG 21:2/MG O-21:3;O | .0010<br>.0034<br>.0034 | [M+H]+<br>[M+Na]+<br>[M+Na]+ | ST 24:0;O4<br>FA 22:0;O2<br>MG O-19:1;O | Petromyzonol<br>13,14-dihydroxy-docosanoic acid<br>1-O-(2R-hydroxy-4Z-nonadecenyl)-sn-glycerol | C24 bile acids, alcohols, and derivatives<br>Hydroxy Fatty Acid<br>Monoradylglycerols- Monoalkylglycerols |
| 411.348 | DG O-22:1/FA 25:1;O2/MG 22:1/MG O-22:2;O | .0035 | [M+Na]+ | MG O-20:0;O | 1-O-(2R-hydroxy-eicosanyl)-sn-glycerol | Monoradylglycerols- Monoalkylglycerols |
| 425.3637 | DG O-23:1/FA 26:1;O2/MG 23:1/MG O-23:2;O | .0036 | [M+Na]+ | MG O-21:0;O | 1-O-(2R-hydroxy-heneicosanyl)-sn-glycerol | Monoradylglycerols- Monoalkylglycerols |
| 453.395 | DG O-25:1/FA 28:1;O2/MG 25:1/MG O-25:2;O | .0012 | [M+H]+ | FA 28:2;O2 | 2-Hydroxy-24-keto-octacosanolide | Fatty esters- Lactones |
| 729.5079 | LPG 34:3/LPG O-34:4;O/PA 37:2;O/PG O-34:3 | .0014<br>.0038 | [M+H]+ | PG O-34:4<br>PG O-32:1 | PG(O-16:0/18:4(6Z,9Z,12Z,15Z))<br>PG(O-16:0/16:1(9Z)) | 1-alkyl,2-acylglycerophosphoglycerols<br>1-alkyl,2-acylglycerophosphoglycerols |
| 731.5236 | LPG 34:2/LPG O-34:3;O/PA 37:1;O/PG O-34:2 | .0015<br>.0039 | [M+H]+ | PG O-34:3<br>PG O-32:0 | PG(O-16:0/18:3(6Z,9Z,12Z))<br>PG(O-16:0/16:0) | 1-alkyl,2-acylglycerophosphoglycerols<br>1-alkyl,2-acylglycerophosphoglycerols |
| 755.5233 | PA 39:3;O/PG O-36:4 | .0012<br>.0036 | [M+H]+ | PG O-36:5<br>PG O-34:2 | PG(O-16:0/20:5(5Z,8Z,11Z,14Z,17Z))<br>PG(O-16:0/18:2(9Z,12Z)) | 1-alkyl,2-acylglycerophosphoglycerols<br>1-alkyl,2-acylglycerophosphoglycerols |
| 757.539 | PA 39:2;O/PG O-36:3 | .0012<br>.0036 | [M+H]+ | PG O-36:4<br>PG O-34:1 | PG(O-16:0/20:4(5Z,8Z,11Z,14Z))<br>PG(O-16:0/18:1(9Z)) | 1-alkyl,2-acylglycerophosphoglycerols<br>1-alkyl,2-acylglycerophosphoglycerols |
| 759.5547 | PA 39:1;O/PG O-36:2 | .0013<br>.0037 | [M+H]+ | PG O-36:3<br>PG O-34:0 | PG(O-16:0/20:3(8Z,11Z,14Z))<br>PG(O-16:0/18:0) | 1-alkyl,2-acylglycerophosphoglycerols<br>1-alkyl,2-acylglycerophosphoglycerols |
| 781.5392 | PA 41:4;O/PG O-38:5 | .0014<br>.0038 | [M+H]+<br>[M+Na]+ | PG O-38:6<br>PG O-36:3 | PG(O16:0/22:6(4Z,7Z,10Z,13Z,16Z,19Z))<br>PG(O-16:0/20:3(8Z,11Z,14Z)) | 1-alkyl,2-acylglycerophosphoglycerols<br>1-alkyl,2-acylglycerophosphoglycerols |

**Table S2**. Masses and potential structures of significantly increased lipids as reported by OrbiSIMS in domestic cat (Felis catus) compared to domestic dog (Canis domesticus familiaris) and Scottish wildcat (Felis silvestris), with classes determined by LIPID MAPS™. Key: † delta scores are an indication of mass tolerance (+/- m/z), set at 0.005 = higher probability that the identification is correct. *No hits achieved at delta 0.005m/z, therefore these are reported at 0.05m/z.

**Figure S5:** Spatial distribution of top 20 lipids exclusive to domestic cat

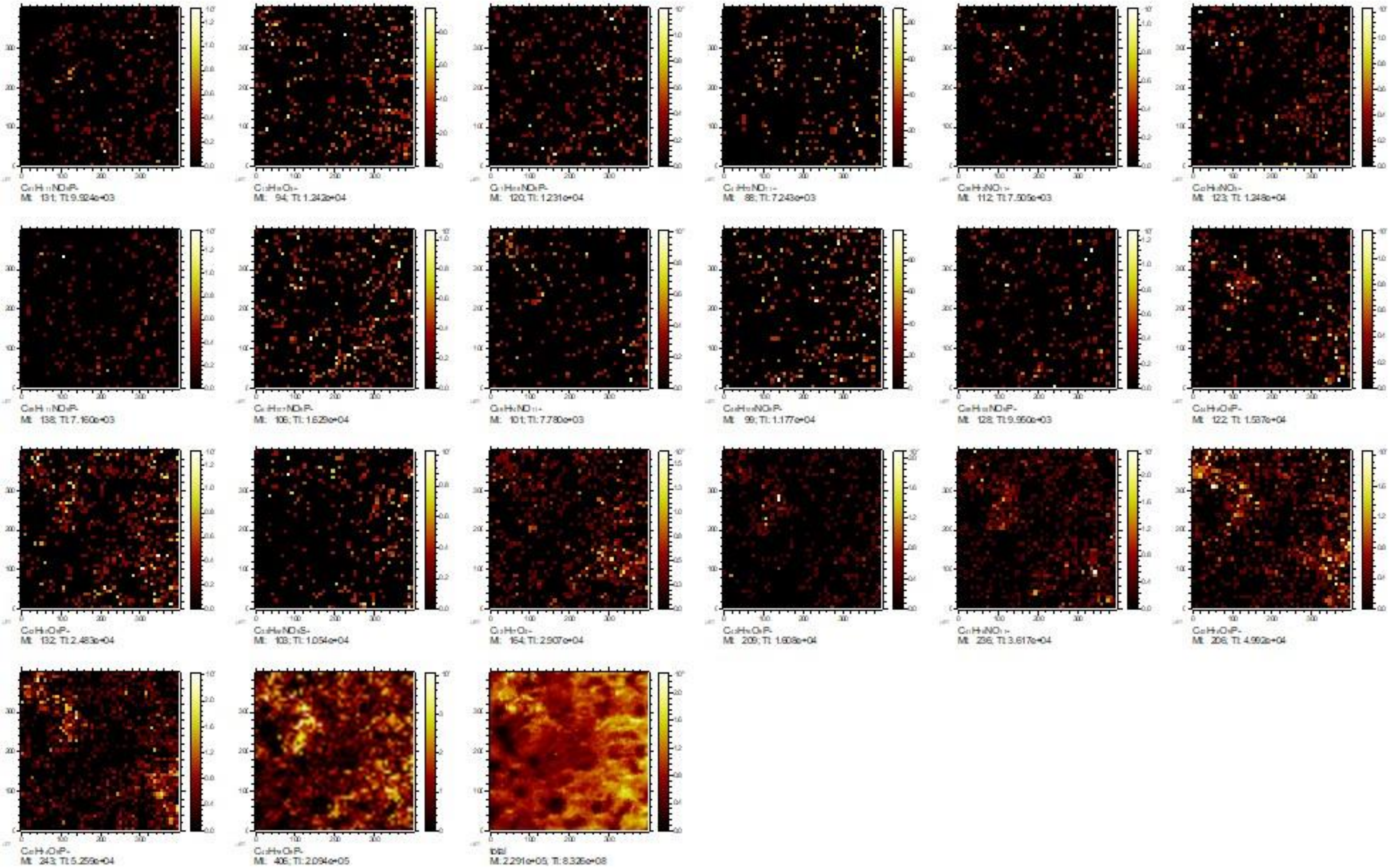

**Figure S5.** Ion images of the top 20 lipids exclusive to domestic cat kidneys (with kidney tissue areas of 400 × 400 µm). Heat map distribution ranges from black through red, yellow and white as intensity increases. The first 20 images are divided by individual lipid intensity. The 21st image is the total intensity of all 20, summated.

**Supplementary Item 8:** LipidMaps assignment of 347 peaks unique to domestic cat (see excel file, n=347 lipids exclusive to DC_OrbiSIMS.xlsx). (rows in green are known MADAGS from Bartz et al, 2007.)
